# Genomic characterization of patient derived xenograft models of multiple myeloma reveals selection of multiple pre-existing high risk clones

**DOI:** 10.64898/2026.10.14.755992

**Authors:** Nathan J. Becker, Enze Liu, Riya Sharma, Parvathi Sudha, Sara Ortega Bertran, Oumaima Jaouadi, Sreejeta Mondal, Nelia Luviano, Maurizio Affer, Kiran K. Mangalaparthi, Akhilesh Pandey, Marios Papadimitriou, Benjamin T. Diamond, David G. Coffey, Abhishek Pandey, James Hoffman, Gil Hevroni, Dickran Kazandjian, Kelvin Lee, Gareth J. Morgan, Faith E. Davies, Attaya Suvannasankha, Aneta Mikulasova, Ola Landgren, Rafat Abonour, Brian A. Walker

**Author notes:** **Corresponding author:** Brian A. Walker, 1550 NW 10^th^ Ave, Suite PAP-511, University of Miami, Miami, FL, USA, 33136.

## Abstract

Multiple myeloma (MM) is a genomically complex malignancy of terminally differentiated plasma cells. Here, we have utilized the SCID-rab patient derived xenograft (PDX) model as a platform to expand scarce primary MM samples across disease stages and major cytogenetic subgroups for comprehensive molecular analysis. The model generated abundant CD138+ cells while retaining key genomic, transcriptomic and proteomic features of corresponding patient tumors. Compared with human myeloma cell lines, PDX cells more closely resembled primary MM in proliferative state and expression of therapeutically relevant targets. Whole genome sequencing (WGS) and single-cell WGS showed that PDXs arose from the selection of pre-existing subclones, including clones harboring high-risk alterations and convergent mechanisms of tumor suppressor inactivation. These findings establish SCID-rab PDX models as a platform for multi-omic profiling, studying clonal evolution, and determining therapeutic response in MM.

## Introduction

Multiple myeloma (MM) is a genomically complex malignancy of terminally differentiated plasma cells and is the second most common hematological cancer.^1^ While patient outcomes have been improving in recent years, MM remains a challenge to treat due to the emergence of high-risk clones that can evolve due to genomic instability and selection.^2,3^ Therefore, having representative models to investigate the diverse and complex molecular mechanisms that drive MM evolution is of critical importance.

Human myeloma cell lines (HMCLs) are derived from advanced stages of disease and do not capture the characteristics of earlier stages such as monoclonal gammopathy of undetermined significance (MGUS), smoldering multiple myeloma (SMM), and newly diagnosed MM,^4,5^ and also lack hyperdiploid (HRD) MM, which accounts for approximately 50% of all diagnoses.^6,7^ Mouse models of MM, including the 5T murine model (5TMM) and the Vk\**MYC* model,^8–10^ have been used to investigate bone disease and treatment responses, but do not possess the classical IgH translocation partners seen in human disease.^9^ Other pre-clinical mouse models that mimic Ig translocations in combination with key driver alterations (*Kras* mutation, *Trp53* deletion, *Bcl2* or *Myc* expression) have also been developed^11^ The advantage of these three systems is that the mice have intact immune systems, allowing interplay between the tumor and the microenvironment to be analyzed.

Patient-derived xenografts (PDXs) enable the expansion of primary human tumor cells in immunodeficient mice, while retaining the principal genomic and transcriptomic characteristics of primary patient samples, making them valuable platforms for studying tumor biology and therapeutic response.^12,13^ The SCID-rab PDX model can support human MM growth within the bone marrow microenvironment of a subcutaneously implanted rabbit bone fragment in immunodeficient mice^14^. allowing the expansion of primary MM samples *in vivo* and unlocking the potential for in-depth multiomic investigation.

Here, we repurpose and refine the SCID-rab model as a platform for expanding primary MM cells from diverse clinical stages and all major cytogenetic subgroups. We evaluate its capacity to generate abundant CD138+ cells and use genomic, transcriptomic, proteomic, and single-cell analyses to determine the extent to which xenografts reflect the corresponding patient tumors. We show that SCID-rab xenografts retain key clinical and molecular characteristics of primary MM while revealing the selective expansion of pre-existing subclones, including populations harboring distinct high-risk clones. These findings establish the SCID-rab model as a complementary, clinically relevant resource for overcoming limitations in patient material and enabling multiomic, functional, and evolutionary studies of MM.

## Results

### SCID-Rab Patient-Derived Xenografts Successfully Expand Patient Samples

We used two variations of the previously described SCID-rab model, employing either NSG or hIL-6-NOG mice as the host strain with CD138^+^ MACS-sorted MM cells injected into an implanted rabbit kit femur fragment (**Figure 1A**).^14^ Initial samples collected from patients with relapse refractory multiple myeloma (RRMM) or plasma cell leukemia (PCL) were injected into NSG SCID-rab mice. Subsequently, as transgenic models became available, samples from newly diagnosed multiple myeloma (NDMM) or smoldering multiple myeloma (SMM) patients were preferentially implanted into hIL-6-NOG mice due to a higher likelihood of dependence on hIL-6.

**Figure 1.**
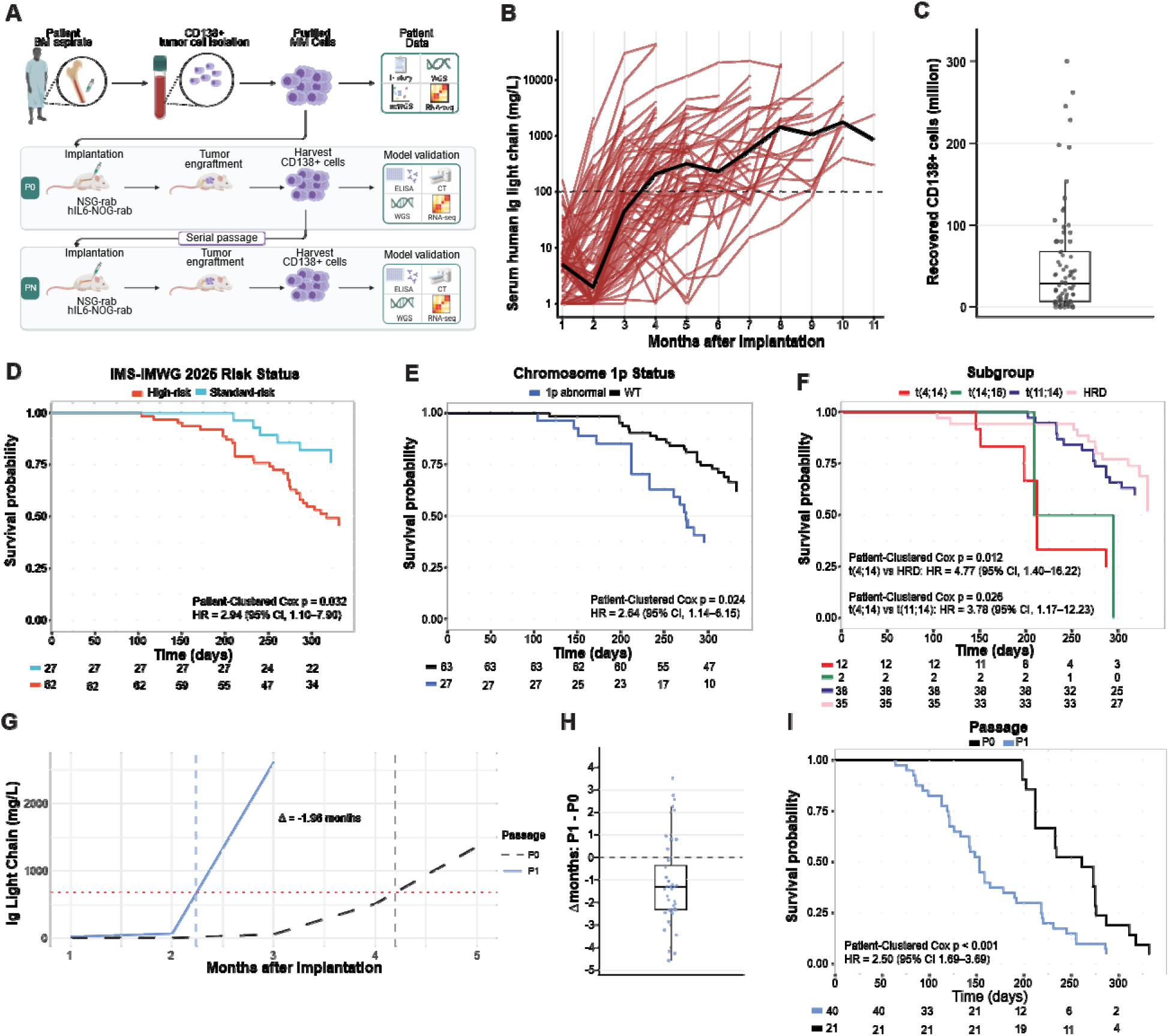
Establishing SCID-Rab PDX models. **A.** The SCID-rab model of multiple myeloma. Patient bone marrow aspirates are CD138+ cell sorted to enrich MM cell populations which are implanted into SCID-rab mice. Cells were collected at the humane endpoint from the rabbit bone and were used in model validation and further passaged. Created with BioRender. **B.** Monthly serum Ig light chain ELISA values for SCID-rab PDX mice from which CD138+ cells were collected. Red lines indicate individual mouse values and the black line indicates the monthly average across all mice. Dotted line represents a derived threshold indicating successful engraftment. **C.** The number of CD138+ MM cells recovered from each SCID-rab mouse (average = 52.1 million, median = 28.2 million). **D.** Survival analysis of PDX based on IMS-IMWG 2025 high risk criteria (*p* = 0.032) **E.** Survival analysis of PDX based on 1p status (*p* = 0.024). **F.** Survival analysis of PDX mice based on cytogenetic subgroup (*p* = 0.012). **G.** P1 mice reached a relative ELISA threshold, defined as 50% of the maximum ELISA value of the matched P0, earlier than their corresponding P0 mice. The black dashed line indicates half P0 max. **H.** Change in half P0 max (Δ) across mice. Crossing times were estimated by linear interpolation between monthly ELISA measurements; for curves that did not cross the threshold, the month of peak ELISA signal was used. **I.** Survival analysis of primary implant (P0) vs. reimplant (P1) mice.

Monthly human immunoglobulin (Ig) light chain ELISA and serum protein electrophoresis (SPEP) were used to monitor engraftment and disease burden, and captured the progressive increase in circulating human Ig light chain (**Figure 1B**, **Extended Data Figure 1A-B**).^14,15^ Successful engraftment, where CD138+ cells were recovered, was highly likely when Ig light chain levels exceeded 100 mg/L, which occurred at a median of 3.5 months post-cell injection. Mice were euthanized at the humane end-point, which occurred at a median of 9.4 months post-cell injection, and the rabbit bone fragment was excised from the mouse flank (**Extended Data Figure 1C-E**). The average number of CD138+ cells collected per SCID-rab mouse was 52.1 million (median = 28.2 million) (**Figure 1C**) and the highest number of cells isolated was 300 million. Endpoint ELISA light chain values were not predictive of CD138+ cell yield from the rabbit bone (**Extended Data Figure 1F**), in line with clinical observations that light chain concentrations and tumor burden are highly variable between patients.^16^

A total of 54 patient samples were injected into 140 SCID-rab PDX mice, with multiple mice injected per patient sample when sufficient primary material was available. Overall, 79.6% of patient samples showed engraftment by ELISA but only 37% yielded PDX cells to sustain models over time (**Supplementary Table 1**). The number of CD138+ cells collected from the rabbit bone of successfully engrafted mice was not dependent on disease stage or cytogenetic subgroup (**Extended Data Figures 1G-I**).

Of the remaining 34 patient samples where PDX cells were not recovered, 23 samples showed sustained progressive Ig light chain levels in mice over time (**Extended Data Figure 1J**), suggesting that MM cells were growing in the mice but were not recovered in the rabbit bone. This was more pronounced in the hIL6-NOG mice compared to the NSG mice (21 vs. 2), suggesting that the presence of hIL6 in the mouse allowed dissemination of the cells across the animal. The remaining mice showed no sustained serum Ig light chain positivity throughout the course of the experiment **(Supplementary Table 1)**.

Gain of *CKS1B* (chromosome 1q) (75.0% versus 37.5%; OR, 5.00; 95% CI, 1.45–17.27; Fisher’s exact *p* = 0.01), and *TP53* abnormalities (60.0% versus 28%; OR, 3.8; 95% CI, 1.18-12.48; Fisher’s exact *p* = 0.04), were associated with successful engraftment indicating high-risk genomic features may be enriched in these models (**Extended Data Figure 2A-B**).

### High-risk markers in patient samples are reflected in PDX survival

Where clinical DNA sequencing data were available (52/54 samples),^17^ high-risk features were determined,^2^ and incorporated into a Cox proportional hazard model, along with mouse strain (NSG or hIL6-NOG), and disease stage (SMM, NDMM, RRMM, sPCL), which identified that IMS-IMWG 2025 high-risk status (HR, 2.94; 95% CI, 1.10-7.90; *p* = 0.032), and *CDKN2C* status (HR, 2.64; 95% CI, 1.14-6.15; *p* = 0.024) were associated with inferior survival (**Figure 1D-E**). Additionally, the t(4;14) subgroup showed inferior survival compared to the HRD (HR, 4.77; 95% CI, 1.40-16.22; *p* = 0.012) and the t(11;14) subgroups (HR, 3.78; 95% CI, 1.17–12.23; *p* = 0.026) (**Figure 1F**). These findings reflect clinical outcomes in MM, where IMS-IMWG 2025 high-risk status, del(*CDKN2C*), and t(4;14) translocations are associated with adverse outcomes.^2,6^

### Engraftment time decreases with subsequent passages

Once PDX models were established (P0), recovered CD138+ cells were reimplanted into an additional round of SCID-rab mice (P1) enabling further expansion of patient-derived MM cells. In paired P0/P1 lineages, P1 mice reached a relative ELISA threshold earlier than their corresponding P0 mice. The mean difference in time to the threshold was -1.14 months, indicating earlier detection and faster growth in the reimplanted PDX mouse (**Figure 1G-H**, **Extended Data Figure 1K-L**). In addition, we observed that P1 mice had significantly reduced survival compared to their matched P0 counterparts (HR 2.50, 95% CI 1.69–3.69, *p* < 0.001) (**Figure 1I**), suggesting that once patient samples are established, subsequent passages feature disease that is detected earlier, thereby reducing the study duration for future experiments.

### PDX mice display patient disease phenotypes

An important clinical characteristic of MM is the development of bone lesions found throughout the skeleton.^16^ In successfully engrafted SCID-rab PDX mice, we observed that the implanted rabbit bone frequently showed loss of rigidity and structural integrity, consistent with extensive local bone destruction facilitating dissociation of the tissue, which aided in the collection of CD138+ MM cells.

Whole-mouse CT imaging was performed on 35 mice following dissection of the rabbit bone implant. Extensive skeletal disease was detected in 12 mice, characterized by multifocal lesions distributed across the skull, ribs and sternum, vertebral column, pelvis, and limbs; which are commonly affected in clinical disease. In one NDMM t(14;16) case, the patient presented with diffuse osseous disease, and the corresponding PDX mouse exhibited the same pattern, with numerous bone lesions throughout the skeleton (**Figure 2A**). Conversely, in a RRMM t(4;14) case, clinical reports recorded minimal bone disease despite multiple rounds of treatment and relapse. This was reflected in the PDX mouse, which showed no significant bone disease in the skeleton (**Figure 2A-B**, **Supplementary Table 2**).

**Figure 2.**
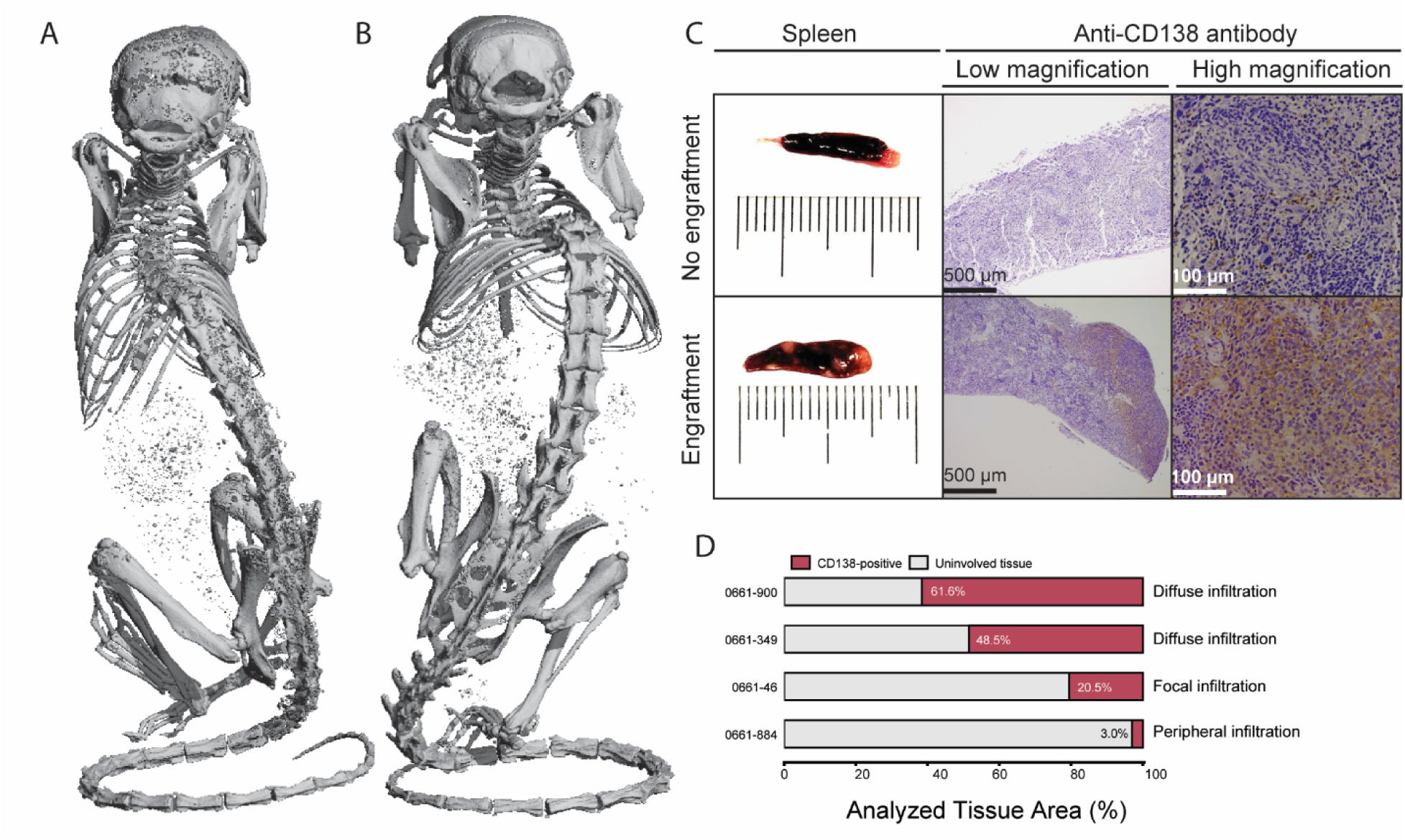
Pathology of PDX mice. **A.** micro-CT scan after removal of the implanted rabbit bone of a SCID-rab mouse derived from a t(14;16) patient showing diffuse osseous disease. **B.** micro-CT scan after removal of the implanted rabbit bone of a SCID-rab mouse derived from a t(4;14) patient showing minimal bone disease. **C.** Spleens from PDX mice with and without successful engraftment and histology at high and low magnification with anti-CD138 DAB staining. **D.** Different patterns and levels of infiltration of MM cells into the spleen were observed.

Splenic infiltration by myeloma cells is a rare event seen in some patients with advanced disease.^18,19^ Enlarged spleens in a subset of SCID-rab mice (4/36) showed extramedullary dissemination by immunohistochemistry (**Figure 2C**), which was either focal (n = 1), diffuse (n = 2), or peripheral (n = 1) (**Figure 2D**, **Extended Data Figure 3**). Together, these findings support the utility of the SCID-rab model to investigate MM-associated bone disease and extramedullary disease.

### SCID-rab PDX mice are a representative model of multiple myeloma patients’ genomic characteristics

A set of SCID-rab PDX mice, derived from 16 patient samples that yielded abundant CD138+ cells, was taken forward for further genomic and transcriptomic characterization. All major cytogenetic subgroups were represented, including t(4;14) (n = 4), t(11;14) (n = 4), t(14;16) (n = 1), t(14;20) (n = 1), hyperdiploid (HRD) (n = 5), and one hyperhaploid^20^ (HRH) patient. Clinical treatment history from diagnosis up to the point of sample collection was recorded for each patient (**Figure 3A**).

**Figure 3.**
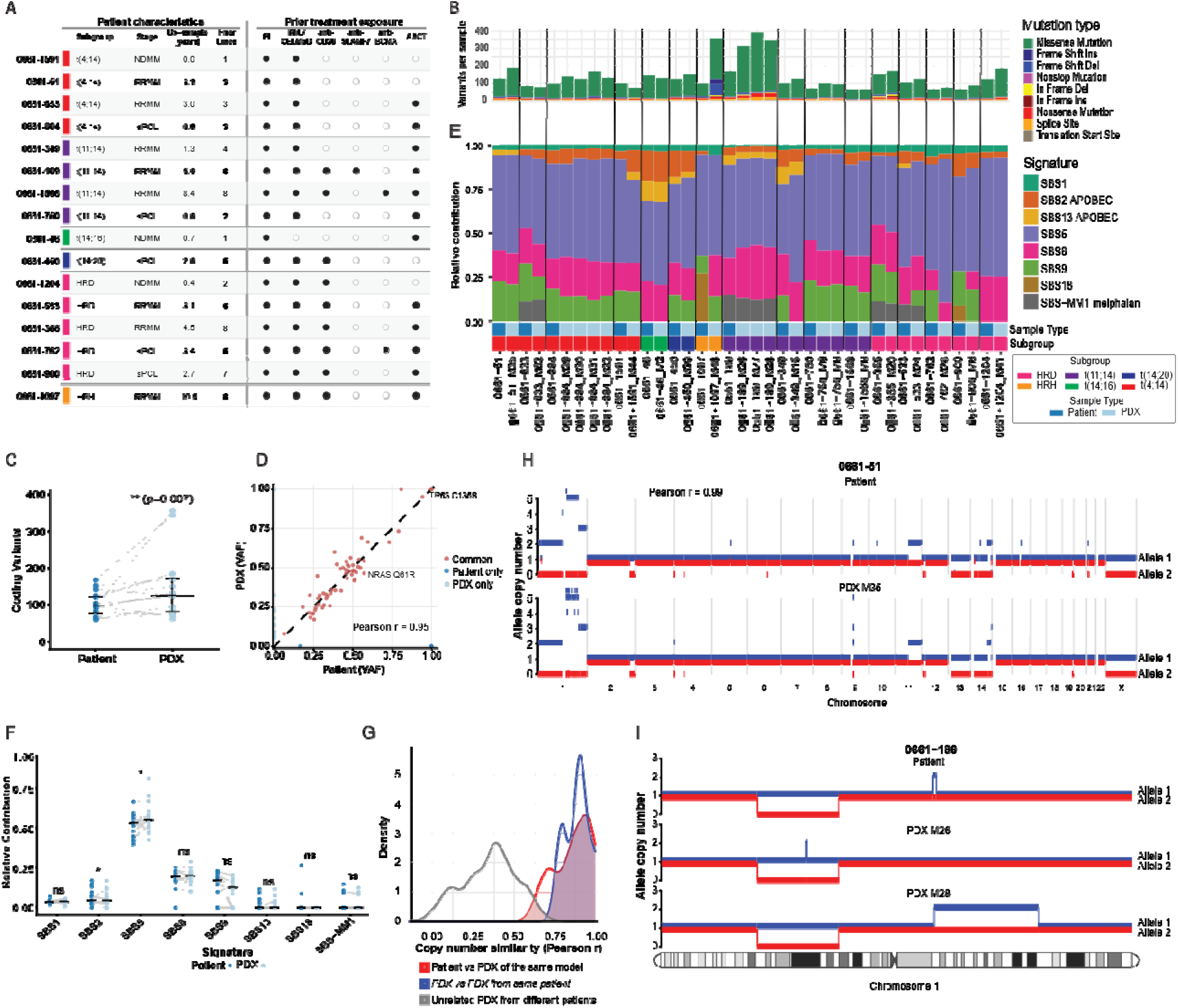
Genomic profiles of patients and their PDX mice. **A.** Clinical characteristics and prior treatment exposure of the 16 patients whose samples underwent multiomic profiling with matched PDXs. Rows represent individual patients and show cytogenetic subgroup, disease stage at sample collection, duration from diagnosis to sample collection, and number of prior treatment lines. **B.** SNVs and Indels detected in each patient and its PDX mice. **C.** The total number of coding variants increases in PDX compared to the primary patient. Gray connecting lines indicate patient and PDX pairs. **D.** Pearson correlation of VAF between patient 0661-762 and its SCID-rab PDX mouse **E.**, MMSIG mutational signatures in patients and their corresponding PDX mice. **F.** Gray connecting lines indicate patient and PDX pairs. **G.** Genome-wide copy number similarity of patients and their PDX mice. **H.** Copy number profile of patient 0661-51 and its SCID-rab mouse shows that the PDX model remains highly similar to the original patient (Pearson *r* = 0.99). **I.** Chromosome 1 CN profile of patient 0661-189 and two matched PDX mice with discordant 1q gain status.

PDX molecular profiles were determined and compared to their matched primary patient samples, indicating an increase in SNVs and Indels in the PDX models (median per patient, 98; median per PDX, 125.5; two-sided paired Wilcoxon signed-rank test, *p* = 0.007; **Figure 3B-C**). Across all PDX samples, there was an accumulation of private mutations, but the rate of accumulation differed across samples and between mice (median = 49; range 12-321). The majority of the private mutations were not in known driver genes of MM and are likely to be passenger mutations. Among variants detected in both paired patient and PDX samples, analysis of variant allele frequency (VAF) showed concordance between PDX models and their primary patient samples (Pearson *r* = 0.58-0.95; **Figure 3D**, **Supplementary Figure 2**). Importantly, matched PDX samples retained many clinically relevant MM driver mutations and CN abnormalities found in the patient (**Figure 4**).

**Figure 4:**
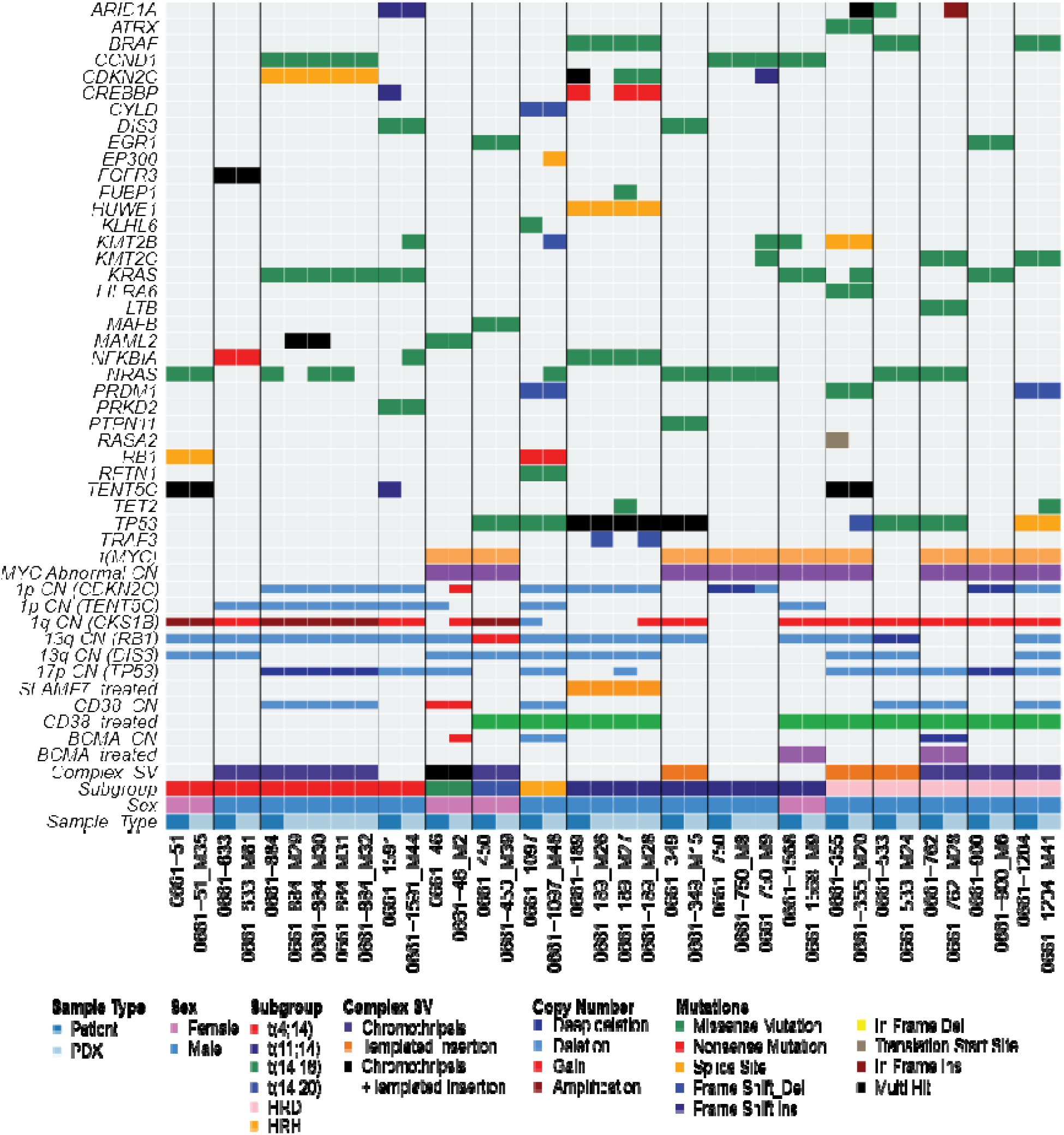
Comparison of patient and derived xenograft common genomic abnormalities.

In addition to SNVs and Indels, mutational signature contributions^21,22^ were compared between matched patient and PDX samples and were largely conserved. A decrease in the SBS2 signature (APOBEC; two-sided paired Wilcoxon signed-rank test, *p* = 0.025), and an increase in the SBS5 (clock-like; *p* = 0.036) were observed (**Figure 3E-F**). The proportional decrease in SBS2 reflected a compositional shift associated with the greater overall SNV burden in PDXs rather than loss of APOBEC-associated mutagenesis. Eleven patients had received high-dose melphalan and autologous stem cell transplantation (HDM-ASCT), and the SBS-MM1 melphalan signature was detected in 4 patients (36%) and was maintained in the PDX model, in line with previous findings that this mutational signature is only present in a subset of patients who received HDM-ASCT.^21^

In a hyperhaploid patient-PDX pair, the PDX exhibited a large increase in genome-wide somatic mutational burden, relative to the matched patient sample, including a substantial accumulation of Indels (**Extended Data Figure 4A**). Examination of the patient and PDX SBS96 mutational spectra further revealed emergence of a prominent CCT>CAT peak in the PDX, which has been previously shown to be associated with disruption of critical DNA mismatch repair (MMR) components *MSH2*, *MSH6* and *MLH1* (**Extended Data Figure 4B**).^23^ Mutational signature fitting identified SBS44, associated with MMR, only in the PDX, (**Extended Data Figure 4C)**.^24^ Interestingly, examination of methylation data showed promoter hypermethylation and gene body hypomethylation of *MSH2*, which was not seen in any other sample (**Extended Data Figure 4D-E**). Two regions containing 6 CpGs in the *MSH2* promoter were identified to be hypermethylated in the PDX. Hypermethylation of this region has been shown to reduce MSH2 expression across other malignancies. When compared to unrelated SCID-rab PDX samples, both CpG regions had significantly higher methylation (Mann-Whitney U p < 0.001). Expression data showed complete loss of *MSH2* expression in the PDX compared to the patient sample, while proteomic analysis similarly showed low MSH2 abundance relative to other PDX models (**Extended Data Figure 4F-G**). Interrogation of the WGS data showed the presence of PDX-acquired Indels at a cumulative frequency of 0.997% in the patient sample, indicating the MMR-deficient clone was present in the original patient sample (**Extended Data Figure 4H)**.

For CN profiles, we found a high degree of similarity between patient and PDX profiles (Pearson *r* = 0.64-0.99), indicating that patient CN landscapes are largely maintained during engraftment (**Figure 3G-H and Supplementary Figure 3**). This represents an important advantage over HMCLs, which often harbor highly abnormal karyotypes, including tetraploidy and CN states rarely observed in primary MM.^25,26^ While the PDX maintained a high degree of similarity, individual PDXs exhibited important mouse-specific differences. In the RRMM patient 0661-189, the patient sample was injected into multiple mice synchronously. Two of the PDX mice had nearly identical CNA profiles, but one had a gain of chromosome 1q that was not detected in the primary sample or the other mice (**Figure 3I**). These findings suggest that the PDX with gain1q arose either through the expansion of a rare, low-abundance subclone or through acquisition of the CNA during growth *in vivo*. PDX-specific differences can be used to better understand MM biology. In this case, the two PDX mice derived from the patient could be used as paired models to investigate the consequences of 1q gain, an important recurrent high-risk feature in MM.

### PDX models retain patient transcriptional features, therapeutic vulnerabilities, and mechanisms of resistance

To determine if SCID-rab PDX cells retain the transcriptional identity of their patients, we compared transcriptional profiles from the paired PDX and patient samples. We found that PDX gene expression was highly correlated with the matched patient sample (Pearson correlation *r* = 0.91-0.98; **Figure 5A and Supplementary Figure 5**). Likewise, we found that PDX models preserved clinically meaningful subtype biology with the expression of subgroup-defining genes and the retention of the patients’ transcriptionally defined classification without gaining a dominant proliferation (PR) signature (**Figure 5B**, **Extended Data Figure 5**).^27^

**Figure 5:**
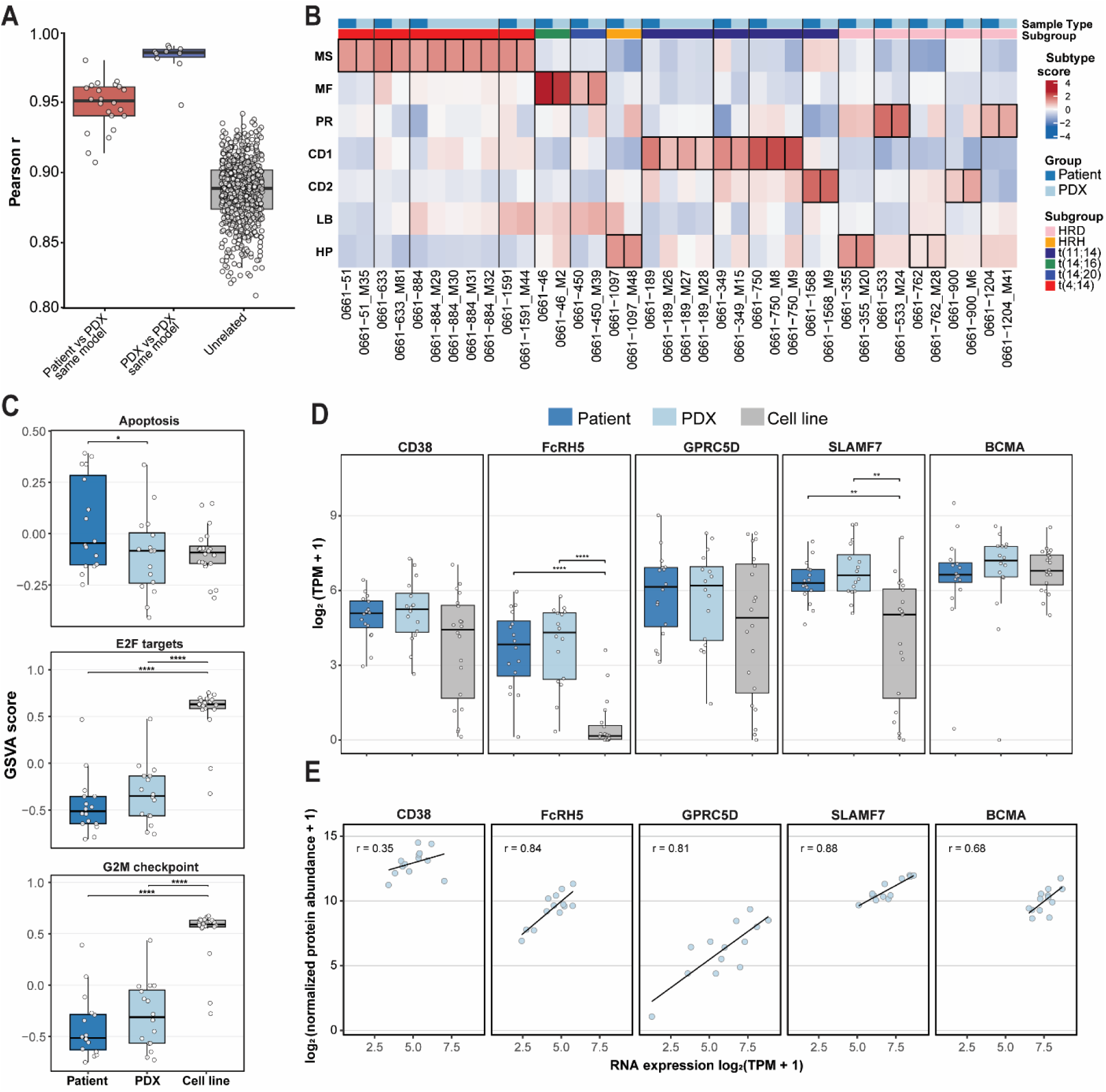
Transcriptomic profiles of patients and their PDX mice. **A.** Pearson correlation of RNA expression between patients and related PDX, related PDX, and unrelated samples. **B.** Transcription subtype scores for patients and their PDX. The subtype with highest score was used to classify the sample. **C.** Patient and matched PDX GSVA scores were compared using two-sided paired Wilcoxon signed-rank tests. HMCL scores were compared with patient or PDX scores using two-sided Wilcoxon rank-sum tests. For patients with multiple linked PDX samples, the median PDX GSVA score was calculated before testing. **D.** Log2(TPM+1) expression of immunotherapy target genes CD38, FcRH5 (FCRL5), GPRC5D, SLAMF7, and BCMA (TNFRSF17) between patients, PDX and cell lines**. E.** Correlation of expression and proteomic data from n=13 SCID-rab PDX for immunotherapy related target proteins.

It has been previously reported that PDX models across cancer types show increased proliferation and decreased apoptosis with increasing passage number relative to the patient sample, suggesting selection for subclones with more aggressive characteristics, which can limit their usefulness as representative models. To investigate this, hallmark G2/M, E2F, and apoptosis gene sets were used to compare patient, PDX, and HMCL samples.^28,29^ HMCLs showed significantly higher expression of hallmark E2F target and G2/M checkpoint genes than PDX and patient samples (unpaired Wilcoxon rank-sum test, both p < 0.001). Consistent with previous findings, PDX samples showed a modest decrease in hallmark apoptosis gene expression relative to patient samples (paired Wilcoxon signed-rank test, p = 0.02). Despite this decrease, SCID-rab PDX samples retained a proliferation-associated expression profile more similar to primary patient samples than did HMCLs, supporting their relevance for modeling patient disease (**Figure 5C**).

### Immunotherapy target gene and protein expression is retained in PDX models

SCID-rab PDXs further preserved clinically relevant immunotherapy target expression (**Figure 5D**). However, when compared to HMCLs (n = 20), patient and PDX samples both significantly expressed higher levels of *FcRH5* (*FCRL5*) and *SLAMF7*. In addition, there was greater variability in the expression of *CD38* and *GPRC5D* in HMCLs compared with patient or PDX samples. This significant difference between PDX cells and HMCLs indicates that PDX myeloma cells may be a more clinically relevant model, as they maintain gene expression critical for testing responses to immunotherapies and support previous findings that HMCLs have highly variable expression of immunotherapy targets.^30^

Proteomic profiling of a subset of the PDX samples (n = 13) further demonstrated that there was a positive correlation between immunotherapy target gene expression and the corresponding protein abundance, although the strength of this relationship varied by target (Pearson correlation *r* = 0.35-0.89) (**Figure 5E**). Preservation of targets at the protein level is an important feature enabling the SCID-rab PDX model to be used as a platform for testing future therapies.

In addition, SCID-rab PDX models may provide a platform for investigating mechanisms of therapeutic escape. Among the 16 patients with paired PDX expression data, 15 had previously received treatment with IMiDs/CELMoDs. Deletions affecting components of the cereblon ubiquitin ligase complex that may confer resistance to IMiD/CELMoD therapy were identified in 10 patients and retained in their PDX mice. Nine patients also received the anti-CD38 immunotherapy daratumumab; of these, four patient samples and their corresponding PDXs had deletion of *CD38*. These findings show the SCID-rab PDXs preserve alterations associated with therapeutic resistance.

### PDX Models Are Derived from Pre-Existing Clones and Identify Multiple High-Risk Subclones Present in the Patient Sample

While SCID-rab PDX mice closely reflect the genetic and transcriptomic profiles of the primary patients from which they were derived, we observed somatic alterations that were either gained during generation or lost during passage in the SCID-rab model. Intratumoral heterogeneity in MM has been widely described, and so we postulated that the differences between the patient sample and the PDX model could be explained by the expansion of pre-existing subclones or ongoing mutagenesis in the cells.^31^ DLP+ scWGS was conducted on a set of six primary patient samples to determine the clonal architecture at the time of implantation into the SCID-rab model.

DLP+ scWGS resolved substantial CN heterogeneity within single-site bone marrow biopsies that was not apparent from bulk WGS alone, identifying a mean of six CN defined subclones per patient. Similar to the previously described MMR-deficient PDX sample, we found that PDX engraftment could be traced to the expansion of pre-existing subclones for all six patients. In one t(11;14) case, bulk-WGS identified biallelic *CDKN2C* inactivation. However, scWGS resolved two distinct *CDKN2C*-inactivated subclones that engrafted in separate PDX mice (**Figure 6A**). One PDX was derived from the dominant patient subclone, which comprised 44% of cells and harbored biallelic deletion of the *CDKN2C* promoter. A separate PDX expanded a rare subclone comprising only 0.5% of cells (**Supplementary Table 3**). This subclone carried a deletion of one *CDKN2C* allele and a R15fs mutation in the remaining allele resulting in biallelic *CDKN2C* inactivation. The R15fs was detected in high-depth targeted panel sequencing at a VAF of 0.03 in the patient, which rose to 0.52 in the PDX and represented the majority of detected transcripts (RNA VAF = 0.76), consistent with deletion of one WT allele and expression of the mutant non-functional allele (**Figure 6B-C**).

**Figure 6:**
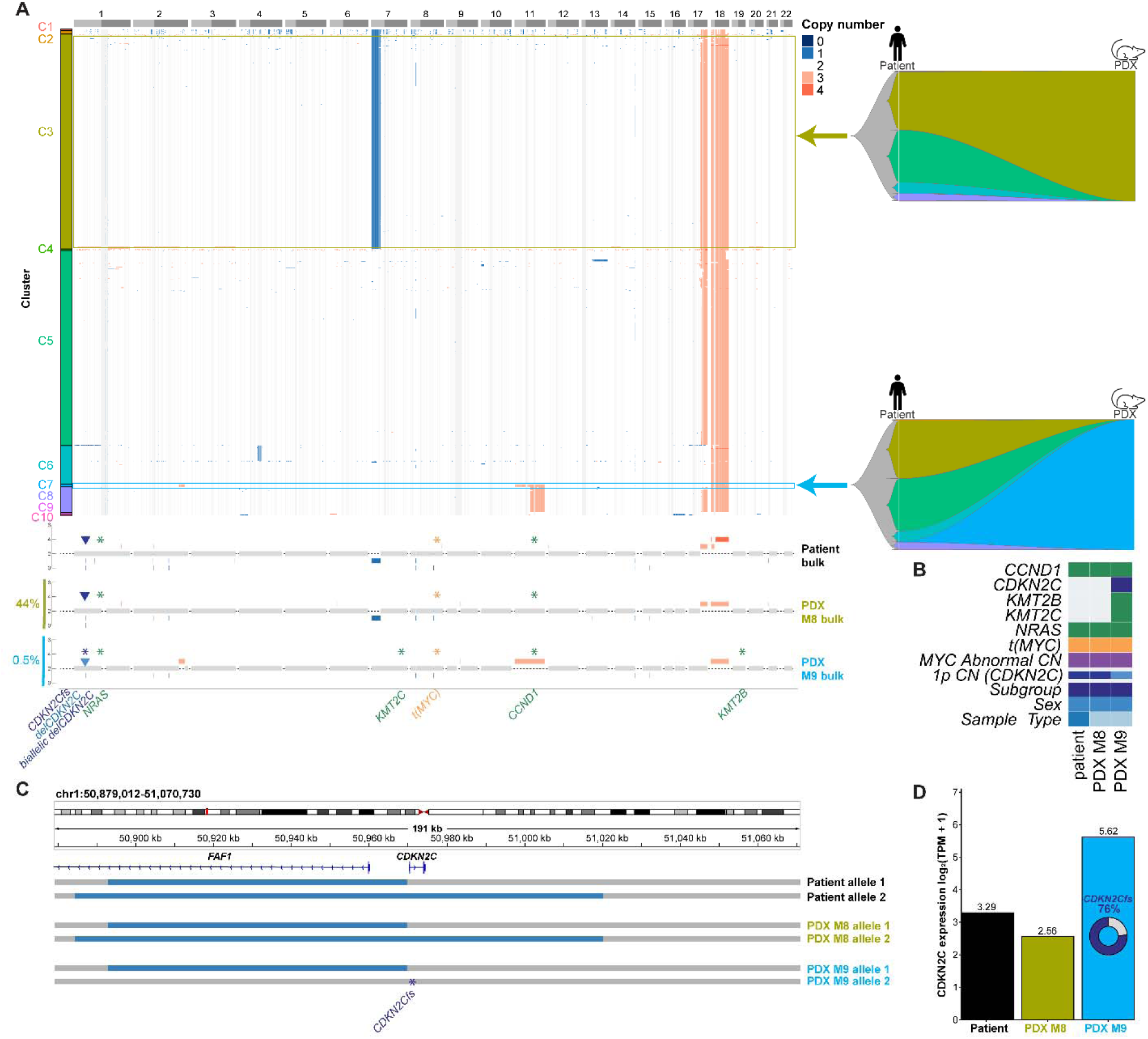
Convergent Mechanisms of biallelic CDKN2C inactivation. **A.** CN defined clones identified by DLP+ scWGS from 967 cells. CN tracks for each sample from bulk WGS are aligned to the scWGS data for comparison. Fishplots represent the clonal composition of the patient sample at the time of implantation and subsequent selection of clones C3 and C7 in the SCID-rab PDX. **B.** MM relevant variants identified by bulk WGS for the patient and corresponding PDXs. **C.** *CDKN2C* localized copy number from bulk-WGS. **D.** RNA expression of *CDKN2C* from PDX M9 highlights the expression of the R15fs allele showing an alternative mechanism of biallelic inactivation.

In a t(4;14) case, bulk WGS identified biallelic deletion of *TP53*. DLP+ scWGS resolved two distinct subclones with biallelic *TP53* inactivation, each of which engrafted in separate mice (**Figure 7A**). The PDXs arose from minor subclones, accounting for 15.2% and 10.6% of the patient population, respectively (**Supplementary Table 3**). The first clone harbored a 17p deletion plus a focal *TP53* deletion, while the second clone featured a focal biallelic *TP53* deletion (**Figure 7B-C**). The focal deletion of *TP53* was not detected by ASCAT-NGS or DLP+ scWGS but was manually detected by inspecting the bulk WGS data (**Supplementary Figure 6**). Within the same patient sample, bulk WGS identified *NRAS* G13V and *KRAS* G12F mutations at a VAF of 0.52 and 0.15, respectively. In three mice, the subclone with the *KRAS* mutation rose to dominance, and in one mouse the *NRAS* and *KRAS* subclones remained at the original frequencies seen in the patient sample (**Figure 7D**).

**Figure 7:**
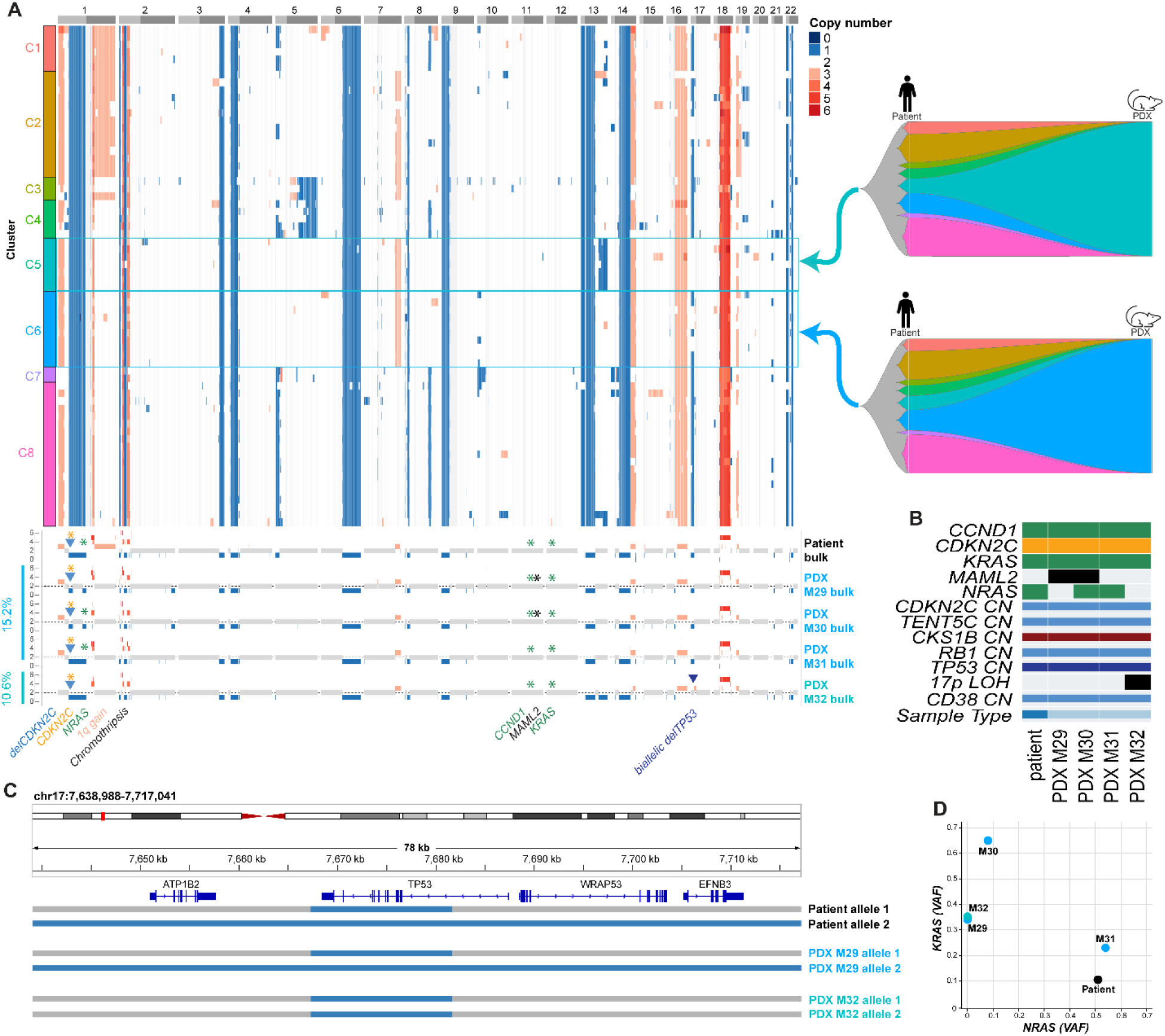
Convergent mechanisms of biallelic inactivation of TP53. **A.** CN defined clones identified by DLP+ scWGS from 66 cells. CN tracks for each sample from bulk-WGS are aligned to the scWGS data for comparison. Fishplots represent the clonal composition of the patient sample and subsequent selection of clones C5 and C6 in the SCID-rab PDX. **B.** MM relevant variants identified by bulk WGS for the patient and corresponding PDXs. **C.** *TP53* localized copy number from bulk-WGS. **D.** VAF of *NRAS* and *KRAS* variants detected in the patient and related PDX mice.

From bulk WGS, we additionally identified a t(11;14) case in which the patient had two *TP53* mutations, R175H and G245R, affecting separate alleles. Two PDX samples generated from this patient retained both *TP53* mutations at frequencies comparable to the primary patient. However, a third PDX showed expansion of a clone harboring a combination of deletions and mutations in alleles. In both instances, the result is biallelic inactivation, indicating ongoing inactivation and alternative mechanisms of generating high-risk biallelic loss of *TP53* (**Extended Data Figure 6**).

Of the six patients analyzed by DLP+ scWGS, four patients contained multiple genomically distinct high-risk subclones, indicating that even in late-stage disease multiple high-risk genetic features can exist in parallel, which may shed light on mechanisms of relapse (**Figures 6-7**, **Extended Data Figures 6-9**). These patterns of selection in the SCID-rab model highlight that high-risk abnormalities are a convergent feature of genetically distinct subclones and that we cannot ignore rare high-risk subclones, since they may have meaningful potential for relapses under selective pressure.

Although PDX engraftment could be traced to the expansion of pre-existing subclones from each patient, we next asked whether engraftment resulted in the emergence of a single dominant population. From the distribution of nonsynonymous somatic mutation VAFs, multiple VAF-defined populations were detected across PDX samples, with a median of 3 inferred clusters in patient samples and 2.5 clusters in PDXs (range, 1–4) (**Supplementary Figure 7**), demonstrating that, despite the expansion of particular subclones during engraftment, most PDXs retained a degree of clonal heterogeneity rather than undergoing a clonal sweep, and suggested that the SCID-rab model preserves multiple tumor populations that may provide a system for investigating ongoing clonal evolution and the effects of subsequent passages.

## Discussion

The ability to expand primary patient MM cells addresses a key barrier to translational research posed by the limited availability of patient samples and their dependence on the bone marrow microenvironment for survival. In this study, we have utilized the SCID-rab model to expand primary patient samples across all major MM cytogenetic subgroups and disease stages, including NDMM, RRMM, and sPCL. The model produced large quantities of viable CD138+ cells for genomic, transcriptomic, and functional analyses while retaining molecular characteristics of the corresponding patient tumors. Cryopreservation and serial passaging extend the utility of these samples, supporting repeated molecular and functional investigations using material derived from the same patient.

The ability to model the biological diversity of MM is particularly important due to the genetic and clinical heterogeneity of MM. Much of our current understanding of MM biology has been derived from HMCLs, or genetically engineered mouse models. Each of these systems remains valuable, but none fully reproduces the clonal complexity, cytogenetic diversity and bone marrow dependence of primary human disease. Integration of patient scWGS data with the SCID-rab PDX model revealed genetically distinct subclones converging on inactivation of the same tumor suppressor genes within individual patients. These findings indicate that even after several lines of therapy, high-risk disease often contains multiple populations with independently acquired high-risk features. Importantly, in several PDX cases, engraftment expanded populations that could be traced back to rare subclones in the corresponding patient samples, making populations poorly resolved by bulk WGS accessible for further characterization. The expansion of these clones demonstrates the possibility that rare high-risk subclones could provide a reservoir for subsequent disease progression. Establishing whether these populations contribute to relapse in patients will require longitudinal clinical sampling in future studies. Additionally, it will be important to implant samples from earlier disease stages with little to no prior treatment to determine if PDX models can predict high-risk clones in patients.

The availability of excess MM PDX cells also creates opportunities for preclinical drug studies. Retention of molecular features of the corresponding patient tumors, together with the ability to cryopreserve and serially propagate these cells, supports repeated evaluation of therapies using material from the same patient. The use of human MM cells also addresses species specific differences that can restrict drug testing in murine tumor models, including differences in cereblon that affect responses to IMiDs/CELMoDs.^32^ Immunotherapy target gene expression is maintained in our PDX model, creating a platform to test immunotherapy drugs.

Our findings establish the SCID-rab model as a platform for expanding patient MM cells while retaining clinically relevant features of disease. Integration with scWGS reveals distinct high-risk populations within individual patients and makes rare subclones accessible for further investigation. These features provide a foundation for studying MM evolution and evaluating therapeutic strategies across all major subgroups in primary human disease.

## Methods

### Patient Samples

Bone marrow aspirates were collected through the Indiana Myeloma Registry with informed consent and institutional approval. All sample collection in this study was reviewed by the Indiana University Human Research Protection Program Institutional Review Board (IRB) and all participants consented in writing to be in this study. Samples were collected under approval 1804208190 (Indiana Myeloma Registry) and this specific study with deidentified samples was deemed exempt by the IUSCCC scientific review committee as the study was retrospective. Additional IRB approval was provided by University of Miami under protocols 20250260 and 20250427.

### Strains of Mice

Two immunodeficient strains of mice were used in this study: NSG (NOD.Cg-Prkdc^scid^ Il2rg^tm1Wjl/SzJ^ Jackson Laboratory) and hIL-6 NOG (NOD.Cg-*Prkdc^scid^ Il2rg^tm1Sug^* Tg(CMV-IL6)1-1Jic/JicTac, Taconic Biosciences). NSG mice were derived from the NOD background and carry the severe combined immunodeficiency (scid) mutation (Prkdc^scid^), combined with a null mutation on the gamma chain of the interleukin-2 receptor (Il2rg) rendering them deficient in functional T cells, B cells, and NK cells. The hIL-6 NOG mice were derived from the standard NOG background (Scid Il2rg^null^) which was modified through the insertion of the human IL-6 transgene. In this mouse strain, hIL-6 is under the control of a CMV promoter and is constitutively expressed. All procedures were approved by the Institutional Animal Care and Use Committee (IACUC) and complied with institutional and federal guidelines for working with immunodeficient animals.

### Generation of SCID-rab PDX mice

SCID-rab mice were generated as previously described, in which the femurs of 3-4 week-old rabbits were cut into two pieces with the proximal and distal ends closed.^14^ Femur sections were inserted subcutaneously through a small incision into either NSG or hIL-6 mice. The bones were left to engraft over the 4 weeks, after which up to 1×10^6^ CD138+ cells were injected into the diaphyseal side of the transplanted femur section. Blood was taken every 4 weeks and serum monitored for MM cell engraftment by immunoglobulin light chain ELISA (Bethyl Labs #NC0102649 kappa and #NC0346121 lambda). Mice were euthanized at the humane end-point or the end of the experiment and the rabbit bone and key organs removed for analysis. Rabbit bones were removed from the mice and crushed to dissociate any CD138+ cells present. Cells were selected for CD138 by MACSsorting (Miltenyi) and viably frozen. Selected cells were passaged by thawing the cells, washing twice in PBS before injecting into a rabbit bone which was engrafted into another mouse.

### µCT Scanning and 3D Reconstruction

Whole mouse carcasses were scanned at 20 μm resolution in a 29.4mm tube using a SCANCO vivaCT-40 system (SCANCO Medical AG, Fabrikweg, Switzerland; 70 peak kilovoltage and 145-µA X-ray source). 3D reconstruction of the µCT images was performed using SCANCO µCT software. Air was the medium of the scan.

### Immunohistochemistry

Formalin-fixed spleen specimens were dehydrated through a graded series of ethanol fixes (70%, 80%, 95%, 100%; two changes of each concentration; 45 minutes per step), cleared in two changes of xylene (45 minutes each), and infiltrated through 3 changes of melted paraffin (∼60° C; 60 minutes each). The specimens were embedded in melted paraffin (melting temperature 56-57 °C) and allowed to harden. Thin sections (4-12 μm) were cut using a rotary microtome and flattened on a heated water bath, floated onto microscope slides and dried.

Formalin-fixed, paraffin-embedded spleen sections were deparaffinized in xylene and rehydrated through graded ethanol series prior to antigen retrieval in citrate buffer (pH 6.0) for 20 min in a 90 °C water bath. Endogenous peroxidase activity was quenched with 3% hydrogen peroxide in methanol, and sections were blocked in 0.5% bovine serum albumin (BSA) in PBS. Sections were incubated overnight at 4 °C with anti–human Syndecan-1 (CD138; Abcam ab130405) at a 1:100 dilution. After washing, a biotinylated anti-rabbit secondary antibody (Vectastain PK-6101) at a 1:200 dilution was applied, followed by avidin–biotin complex incubation. Signals were developed using 3,3′-diaminobenzidine (DAB), and sections were counterstained with Gill’s hematoxylin. Slides were dehydrated, cleared in xylene, and mounted with a xylene-based mounting medium. Hematoxylin and eosin (H&E) staining was performed on parallel sections for histopathological assessment.

Images of DAB stained spleen sections were analyzed using QuPath version 0.7.0. The spleen tissue was manually outlined to exclude non-tissue regions. DAB stained areas (CD138 infiltrated) were identified using the train pixel classifier function and the artificial neural network classifier. Representative regions were manually annotated as CD138 infiltrated tissue or uninvolved splenic tissue, and the classifier was refined using live prediction until the classification agreed visually with the corresponding histological image. Due to staining intensity and tissue appearance differences among the positive samples, a separate pixel classifier was trained and applied to each CD138 infiltrated spleen section. The areas classified as CD138 infiltration and uninvolved tissue were measured within the annotated spleen region. The level of CD138 infiltration was calculated as the area classified as CD138 infiltration divided by the total tissue area and expressed as a percentage.

### Human Myeloma Cell Lines

HMCLs U266, JJN3, ARP1, LP1, L363, MM1-S, KMS27, PCM6, H929, CAG, KMM1, KMS28, AMO1, KMS26, RPMI-8266, KMS12BM, Sachi, KMS12PE, MM1-R, KMS11 were cultivated in cell culture flasks at 37 °C and 5% CO_2_. Growth medium was RPMI with 10% fetal bovine serum (FBS) (Corning, #35-011-CV). For PCM6, growth medium was McCoy’s with 10% FBS and supplemented with 30U/ml hIL-6 (Gibco, #PHC0064).

### Whole Genome Sequencing

Short-read (Illumina) whole genome sequencing was obtained for patient CD138+ cells, matched control saliva or peripheral blood, and PDX cells generated from the patient. For patient and PDX tumor samples, bone marrow aspirates underwent red cell lysis and white blood cells were separated with Ficoll. Tumor cells were selected using CD138+ magnetic beads (Miltenyi) and viably frozen in fetal bovine serum plus 10% DMSO. DNA and RNA were simultaneously extracted using the AllPrep DNA/RNA/miRNA Universal kit (Qiagen). For saliva, samples were obtained in OG-500 tubes (Oragene). Peripheral blood was collected in an EDTA tube and white cells were separated by Ficoll. DNA was extracted by magnetic purification using a Chemagic 360 (Perkin Elmer).

Samples were sequenced with an average coverage of 65.5X for tumor samples and 33.2X for non-tumor controls. The data were processed using the MGP1000 pipeline where translocations were detected using Manta, SNVs and Indels using Strelka2, and copy number abnormalities using ASCAT-NGS.^33–36^ For patients with multiple PDXs, the median variant count was used as the PDX value for the paired analysis. Mutation signatures were determined using all variants and mmsig adapted for hg38.^37^

### DNA Methylation

Methylation data was derived from long-read WGS data (PacBio). Genome-wide CpG methylation was identified using Primrose.^38^ Methylation profiles were established using Pb-CpG-toolkit (https://github.com/PacificBiosciences/pb-cpg-tools) where >50% probability was required to identify a methylated CpG site in a read and ≥4 reads were required.

### Targeted Panel Sequencing

DNA from both CD138+ cells and non-tumor cells were quantified by Qubit (ThermoFisher) and 100 ng DNA used to prepare a sequencing library with unique dual indices (HyperPlus, KAPA Biosystems). Libraries were pooled and hybridized to the previously published Myeloma Genome Project targeted panel (version 22).^39^ Pooled libraries were sequenced using 150 bp paired end reads to a median depth of 426X (range 53-1313X) for tumor samples on a NovaSeq 6000 (Illumina). Sequencing data were aligned to the human genome (hg38) and analyzed using the previously published pipeline.^39^ Translocations were detected by Manta, copy number by CNVkit, and mutations by Strelka2.^33,34^

### RNA-Sequencing

The stranded total RNA prep with Ribo-zero plus (Illumina) was used to generate libraries from 100 ng RNA. Bulk RNA-sequencing was conducted on HMCLs, patient, and PDX tumor samples. Reads were aligned to the hg38 reference genome with gene annotation from Gencode database (V35) using STAR.^40^ Transcript-level expression was measured by transcript-per-million (TPM) using Salmon (Quasi-mode mapping, ‘validate map’ mode), aggregated up to gene-level expression (raw counts and transcript per million [TPM]), and converted to log_2_(TPM+1) to stabilize variance.^28,29^ Gene expression signatures were retrieved from the Molecular Signature Database.^28,29^

### Proteomics

Proteins were extracted from CD138+ PDX cells and quantified as previously described.^41^ Briefly, proteins were extracted, digested with LysC and trypsin, and processed for mass spectrometry using the established workflow. Protein identification and quantification were conducted according to the previously reported analytical pipeline.

### Single-Cell Whole Genome Sequencing

Direct Library Preparation + (DLP+) single-cell whole genome sequencing was conducted on six cryopreserved CD138+ primary patient samples at the New York Genome Center with an average coverage of 0.05X per cell.^42^ Single cells with <250,000 reads or a quality score of less than 0.75 were discarded. HMMcopy BAM files and segmented copy number data were used to infer the subclonal composition of each sample.^43^ Copy number segments were mapped onto non-overlapping 250kb genomic bins. Copy number states were weighted to emphasize CNAs spanning larger genomic regions to elevate real biological changes over potential noise. ENCODE blacklist regions or regions containing technical artifacts were excluded. Copy number defined subclones were identified using Ward.D2 hierarchical clustering based on Pearson correlation. The number of clusters was selected from k::J=::J2-10 using average silhouette width, which measures similarity within clusters and separation between clusters. Clusters represented by fewer than three cells were combined with the nearest neighboring cluster.

### Assessment of PDX Sample Intratumoral Heterogeneity

To assess residual clonal heterogeneity in SCID-rab PDX models, the distribution of nonsynonymous somatic mutation variant allele frequencies (VAFs) was evaluated within each sample using *inferHeterogeneity* in maftools.^44^

### Data Availability

Raw sequencing data have been submitted to dbGaP under accession number phs004479.v1.p1.

## Acknowledgements

The Indiana Myeloma Registry is funded in part by support from the Indiana University Precision Health Initiative, Miles for Myeloma, the Harry and Edith Gladstein Chair, and the Omar Barham Fighting Cancer Fund. Computational infrastructure at Indiana University (Indianapolis, IN) was funded in part by Lilly Endowment Inc. through the Indiana University Pervasive Technology Institute.

B.A.W. is funded by Blood Cancer United (grant 8049-25), U.S. Department of Defense (CA230482), The International Myeloma Society, and the Daniel and Lori Efroymson Chair (Indiana University).

A.M. is supported by the Medical Research Council [UKRI328].

Additional support was provided by the Simon Comprehensive Cancer Center support grant P30 CA082709 and the Sylvester Comprehensive Cancer Center support grant P30 CA240139. These Cancer Center grants provided support to the *In vivo* therapeutics core (Indiana University), histology core (Indiana University), Center for Medical Genomics (Indiana University), and the Oncogenomics Shared Resource (University of Miami). We pay particular thanks to Karen Pollock and Tony Sinn and their team in the *In vivo* therapeutics core at Indiana University.

