## Extended Data for "Genomic characterization of patient derived xenograft models of multiple myeloma reveals selection of multiple pre-existing high risk clones"

**Running title: PDX models in multiple myeloma**

**Authors**: Nathan J. Becker[1], Enze Liu[1], Riya Sharma[2], Parvathi Sudha[2], Sara Ortega Bertran[1], Oumaima Jaouadi[2], Sreejeta Mondal[1], Nelia Luviano[1], Maurizio Affer[1], Kiran K. Mangalaparthi[3], Akhilesh Pandey[3], Marios Papadimitriou[1], Benjamin T. Diamond[1], David Coffey[1], Abhishek Pandey[1], James Hoffman[1], Gil Hevroni[1], Dickran Kazandjian[1], Kelvin Lee[2], Gareth J Morgan[4], Faith E Davies[4], Attaya Suvannasankha[2], Aneta Mikulasova[5], Ola Landgren[1], Rafat Abonour[1], Brian A. Walker[1]

### Extended Data Figures

**Extended Data Figure 1. PDX model characteristics. A.** SPEP of the PDX 0661-1568 M9 shows an increase in detection of M-spike in PDX serum over time. **B.** ELISA values for 0661-1568 **C.** SCID-rab PDX mouse with successful engraftment of patient CD138+ cells. **D.** Excised rabbit kit femur fragment encapsulated in MM tissue. **E.** Successful engraftment encapsulates the implanted rabbit femur fragment leading to an increase in size when compared to rabbit femurs without successful engraftment $\boldsymbol{p < 0.001}$. **F.** Endpoint serum light chain levels detected by ELISA and the number of CD138+ cells recovered showed no correlation. Points represent individual PDX mice. The black line indicates a linear fit. **G-I.** There was no significant difference in the number of cells recovered between cytogenetic subgroup, mouse strain, or passage number. **J.** Monthly serum light chain values for PDX samples that did not yield recoverable CD138+ cells. A progressive ELISA rise was defined as a peak light-chain level of at least 100 mg/L, a minimum threefold increase over the first measured value, at least two positive timepoints, and positivity at consecutive timepoints. The remaining mice showed low or stable light-chain levels, an isolated positive measurement, or positivity that did not meet the criteria for sustained progression. **K.** Monthly ELISA values from paired P0 and P1 mice. **L.** CD138+ MM cells Collected from paired P0 and P1 mice. Connecting lines indicate P0-P1 pairs.

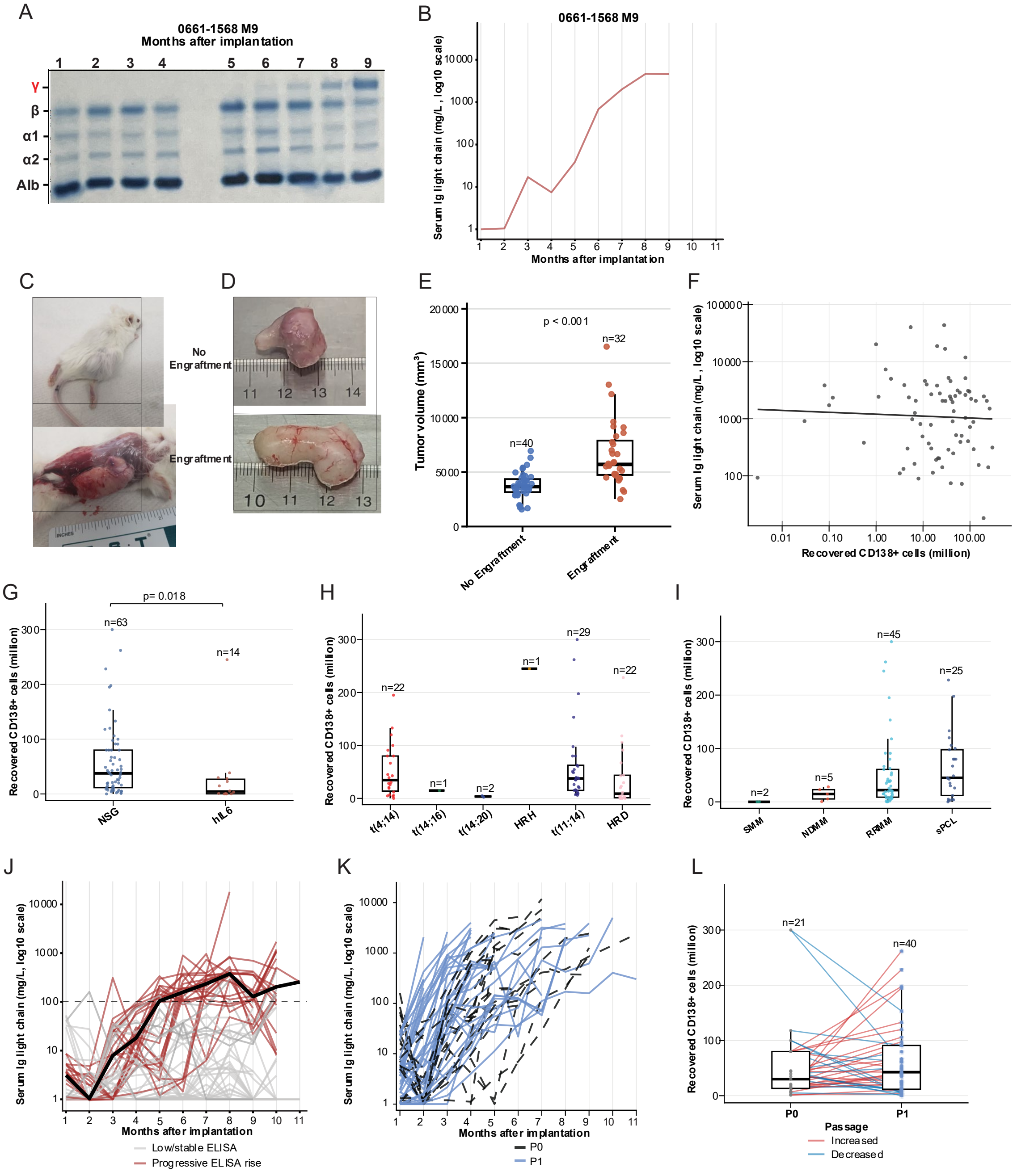
A
0661-1568 M9
Months after implantation
1 2 3 4 5 6 7 8 9
γ
β
α1
α2
Alb
B
0661-1568 M9
Serum Ig light chain (mg/L, log10 scale)
Months after implantation
C
D
No Engraftment
Engraftment
E
Tumor volume (mm³)
p < 0.001
n=40
n=32
No Engraftment
Engraftment
F
Serum Ig light chain (mg/L, log10 scale)
Recovered CD138+ cells (million)
G
Recovered CD138+ cells (million)
p= 0.018
n=63
n=14
NSG
hIL6
H
Recovered CD138+ cells (million)
n=22
n=1
n=2
n=1
n=29
n=22
t(4;14)
t(14;16)
t(14;20)
HRH
t(11;14)
HRD
I
Recovered CD138+ cells (million)
n=2
n=5
n=45
n=25
SMM
NDMM
RRMM
sPCL
J
Serum Ig light chain (mg/L, log10 scale)
Months after implantation
Low/stable ELISA
Progressive ELISA rise
K
Serum Ig light chain (mg/L, log10 scale)
Months after implantation
P0
P1
L
Recovered CD138+ cells (million)
n=21
n=40
P0
P1
Passage
Increased
Decreased

**Extended Data Figure 2. Analysis of patient genetic characteristics based on engraftment vs. failure to engraft in the SCID-rab model. A.** SNVs and Indels for patient samples implanted into the SCID-rab model divided by engraftment status. **B.** Odds ratio for MM relevant variants in engraftment vs. failure.

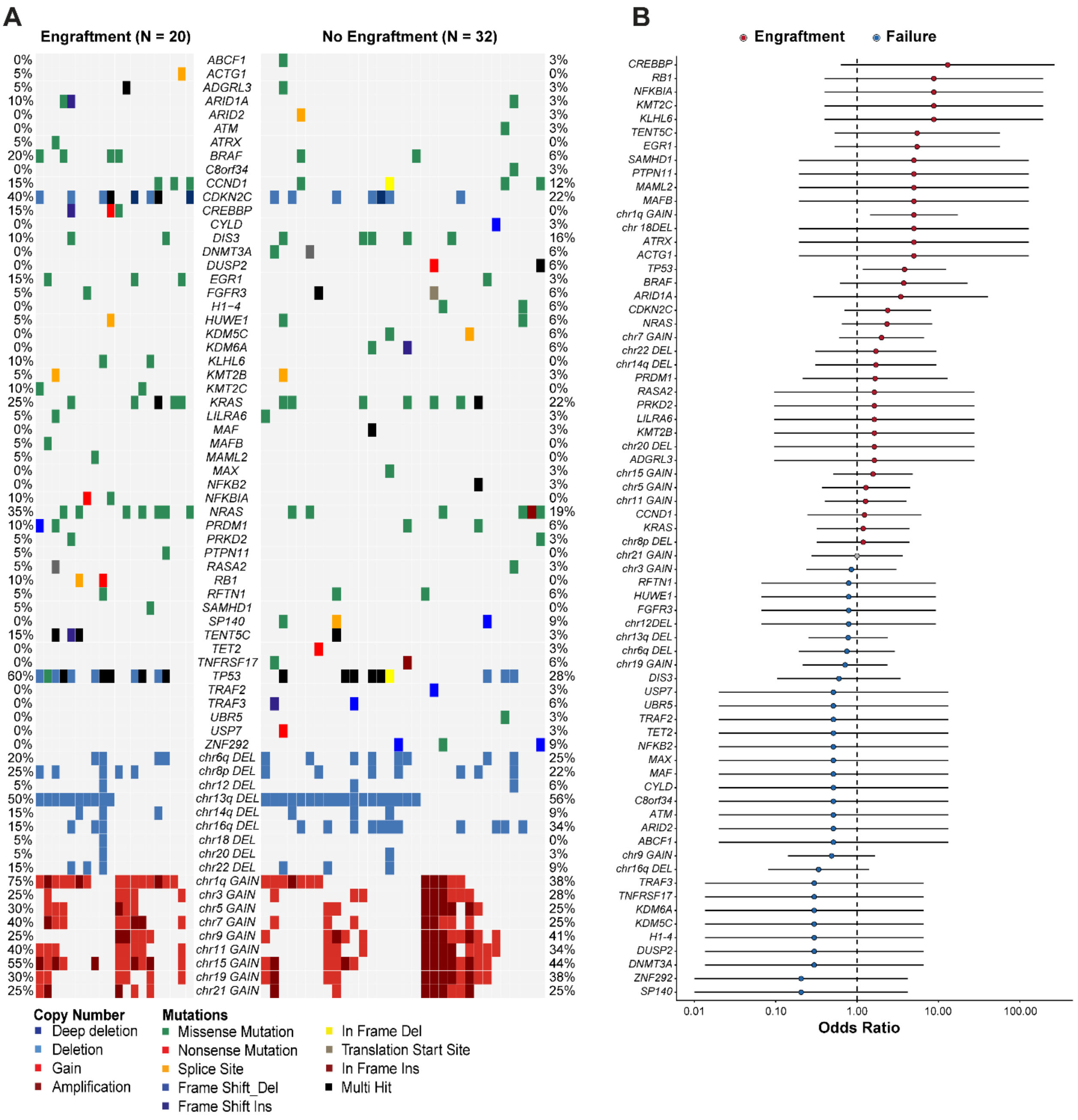

### Extended Data Figure 3. CD138+ infiltration of SCID-rab spleens

**CD138+ spleen sections from SCID-rab PDX mice visualized at low and high magnification.**

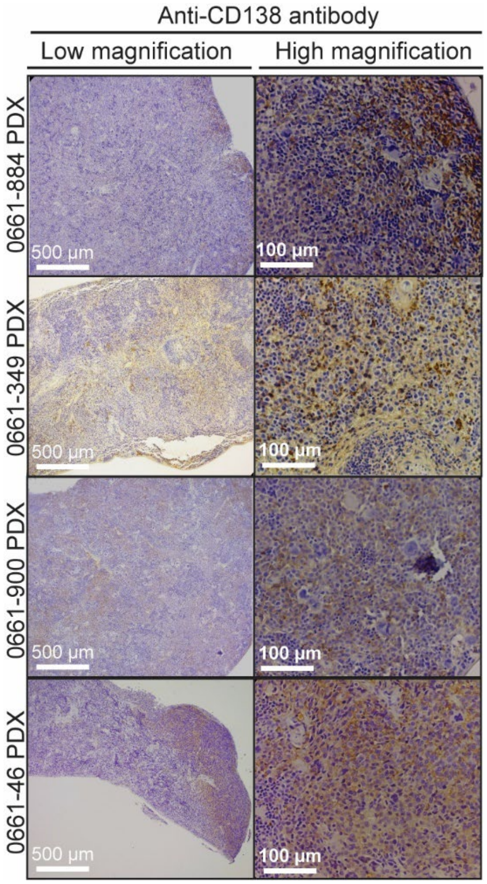

**Extended Data Figure 4. Mismatch repair deficiency identified in a hyperhaploid PDX model.**

**A.** Mutational burden of a matched patient and PDX pair showing increased occurrence of indels in the PDX. **B.** SBS96 spectra of the matched patient and PDX samples showing a CCT>CAT peak in the PDX. **C.** Mutational signature contributions fitted to the patient and PDX samples indicating an increase in SBS44 MMR-deficiency signature. **D.** *MSH2* gene body hypomethylation in the PDX sample with MMR-deficiency compared to other PDX models. **E.** *MSH2* promoter CpGs in the MMR-deficient PDX were hypermethylated. All other PDX models with methylation data are shown for comparison. **F.** Expression data for patient samples and related PDX models show loss of *MSH2* expression in the affected PDX. **G.** *MSH2* protein expression among available PDX samples (n=13) with available data. **H.** Representative IGV screenshot showing a clonal Indel identified in the PDX that was also present at a low VAF in the patient sample BAM file. Unique Indels identified in the PDX (n=179,036) were queried in the patient BAM file where 130,987/13,139,996 reads (0.997%) contained the Indel identified in the PDX.

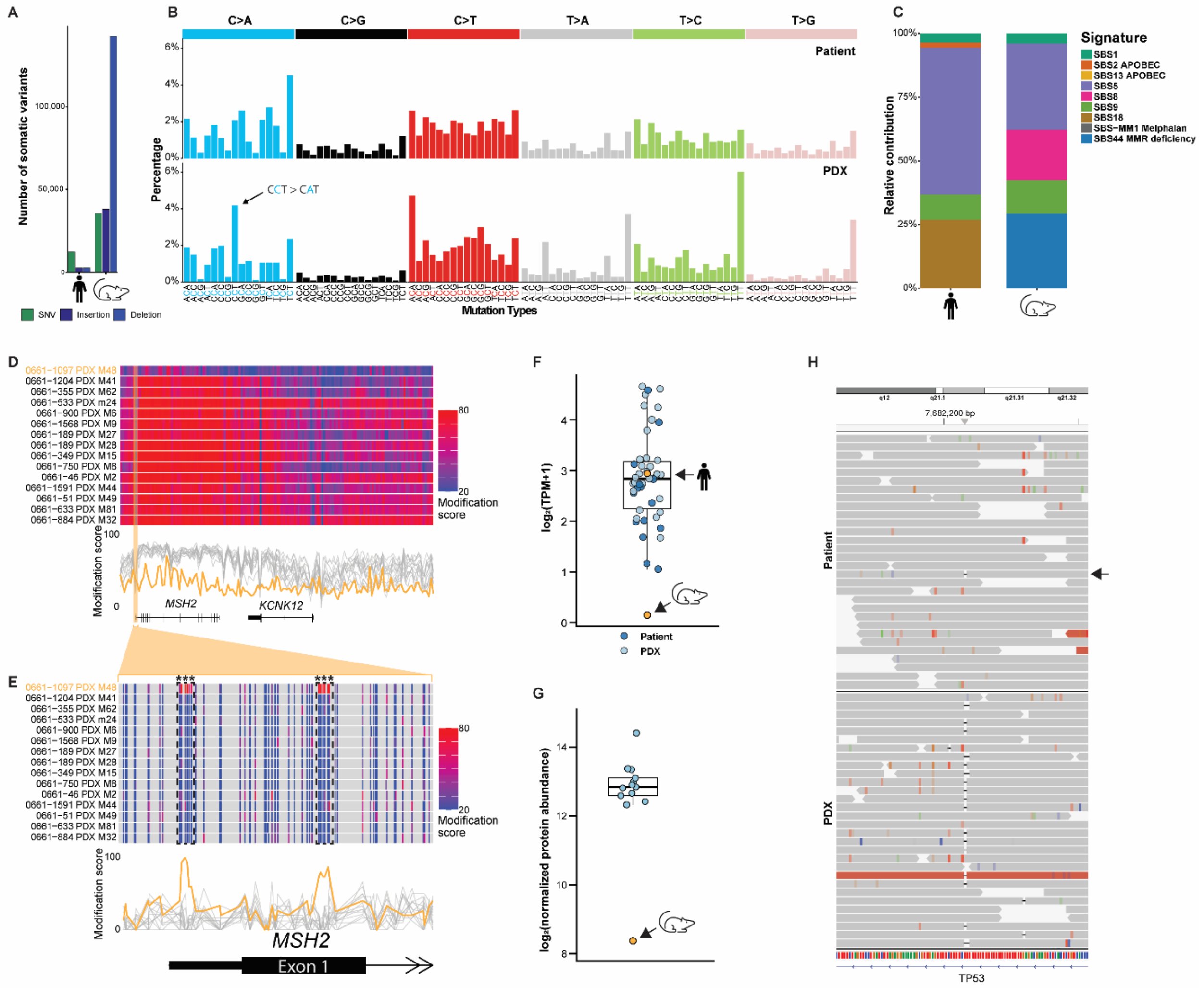
A
Number of somatic variants
SNV
Insertion
Deletion
B
C>A
C>G
C>T
T>A
T>C
T>G
Patient
PDX
Percentage
CCT > CAT
Mutation Types
C
Relative contribution
Signature
SBS1
SBS2 APOBEC
SBS13 APOBEC
SBS5
SBS8
SBS9
SBS18
SBS-MM1 Melphalan
SBS44 MMR deficiency
D
0661-1097 PDX M48
0661-1204 PDX M41
0661-355 PDX M62
0661-533 PDX m24
0661-900 PDX M6
0661-1568 PDX M9
0661-189 PDX M27
0661-189 PDX M28
0661-349 PDX M15
0661-750 PDX M8
0661-46 PDX M2
0661-1591 PDX M44
0661-51 PDX M49
0661-633 PDX M81
0661-884 PDX M32
Modification score
MSH2
KCNK12
E
MSH2
Exon 1
F
log2(TPM+1)
Patient
PDX
G
log2(normalized protein abundance)
H
7,682,200 bp
Patient
PDX
TP53

**Extended Data Figure 5. Expression of subtype defining genes in patient and PDX**

Expression of subgroup defining genes gray lines indicate a paired patient and PDX samples.

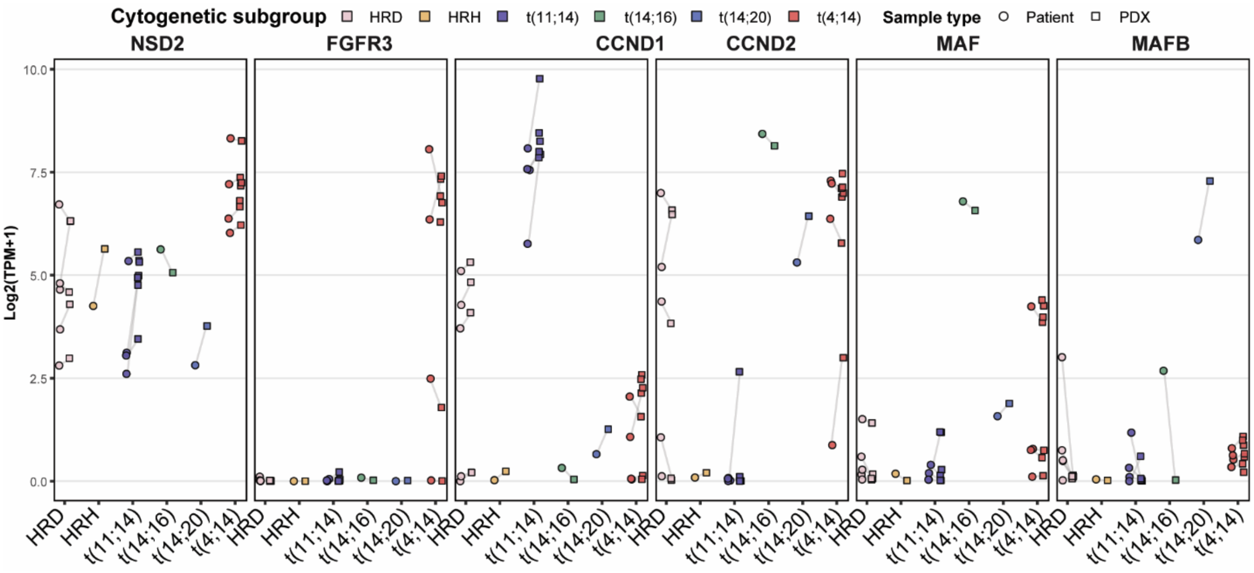

**Extended Data Figure 6. scWGS DLP+ sequencing 0661-189, a t(11;14) sample.**

**A.** CN defined clones identified by DLP+ scWGS from 659 cells. CN tracks for each sample from bulk WGS are aligned to the scWGS data for comparison. Fishplots represent the clonal composition of the patient sample at the time of implantation and subsequent selection in the SCID-rab PDX. **B.** MM relevant variants identified by bulk WGS for the patient and corresponding PDXs. **C.** VAF of *TP53* variants detected in the patient and related PDX mice.

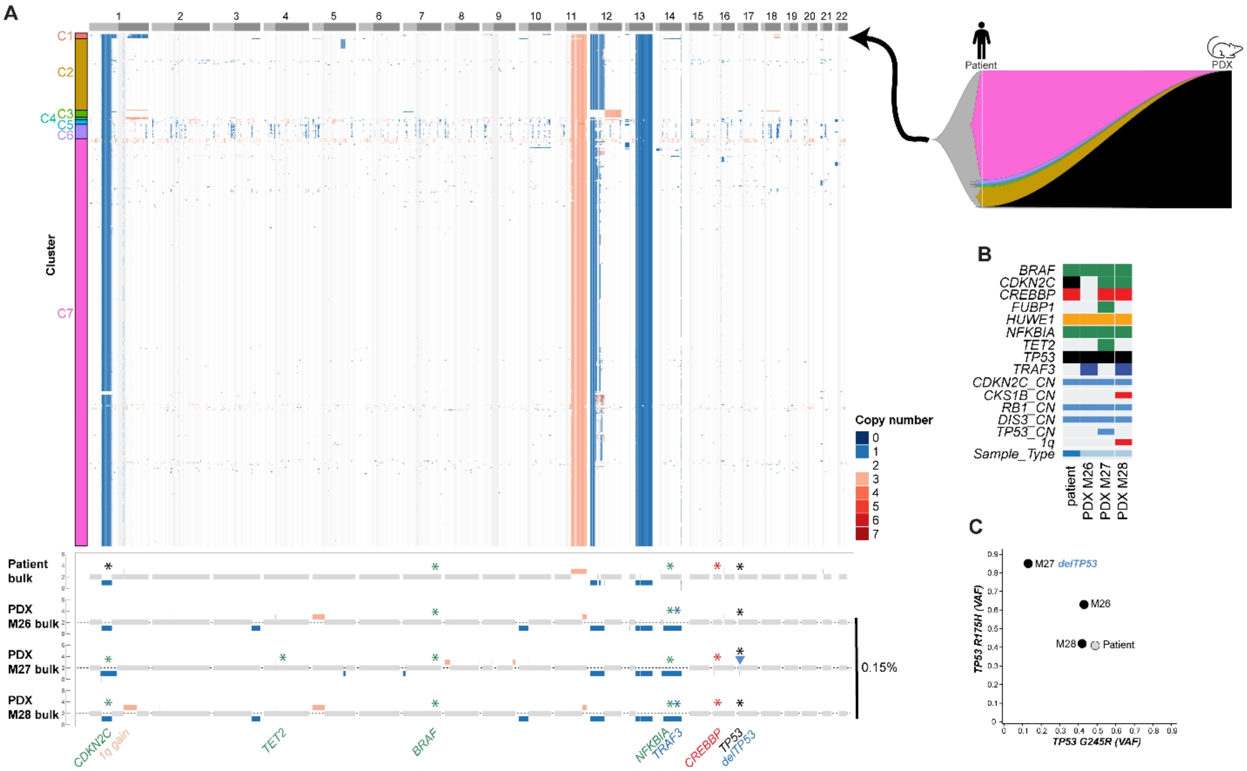

**Extended Data Figure 7. scWGS DLP+ sequencing 0661-633, a t(4;14) sample.**

**A.** CN defined clones identified by DLP+ scWGS from 53 cells. CN tracks for each sample from bulk WGS are aligned to the scWGS data for comparison. Seven main clusters were identified. Fishplots represent the clonal composition of the patient sample at the time of implantation and subsequent selection of clone C2 in the SCID-rab PDX. **B.** MM relevant variants identified by bulk WGS for the patient and corresponding PDXs.

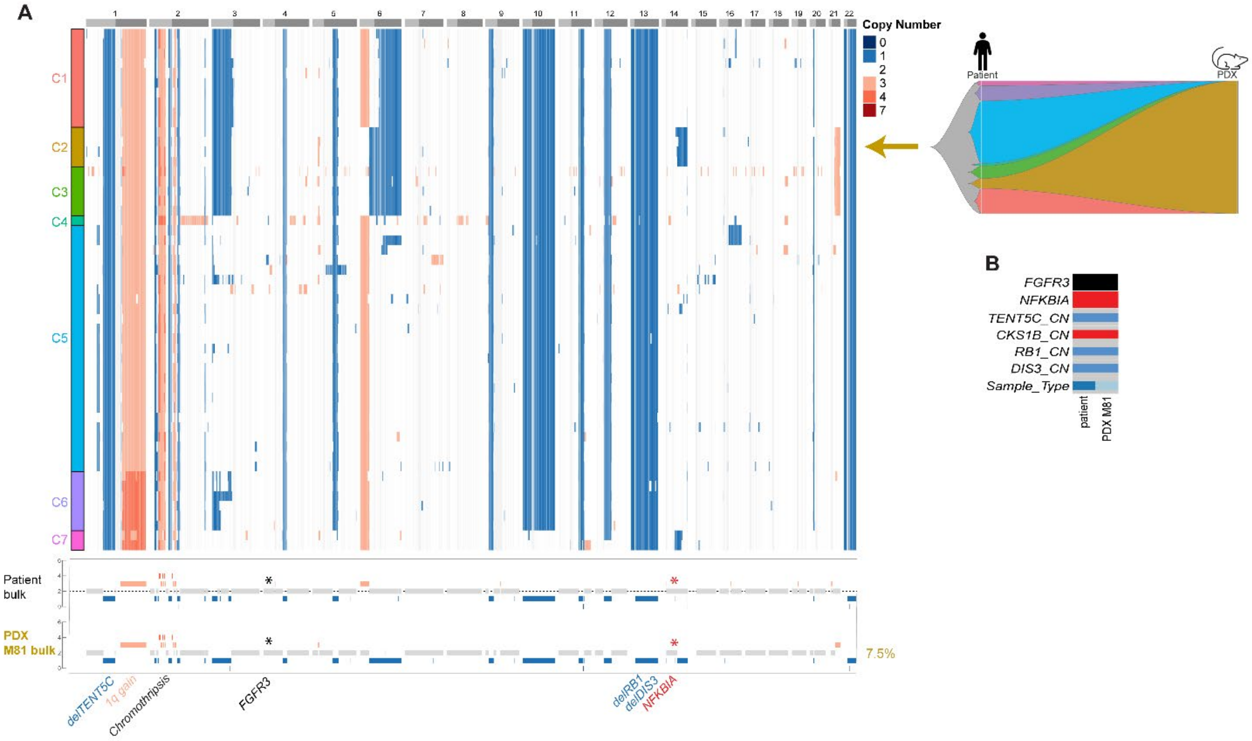

**Extended Data Figure 8. scWGS DLP+ sequencing 0661-349, a t(11;14) sample.**

**A.** CN defined clones identified by DLP+ scWGS from 624 cells. CN tracks for each sample from bulk WGS are aligned to the scWGS data for comparison. Fishplots represent the clonal composition of the patient sample at the time of implantation and subsequent selection of clone C3 in the SCID-rab PDX. **B.** MM relevant variants identified by bulk WGS for the patient and corresponding PDXs.

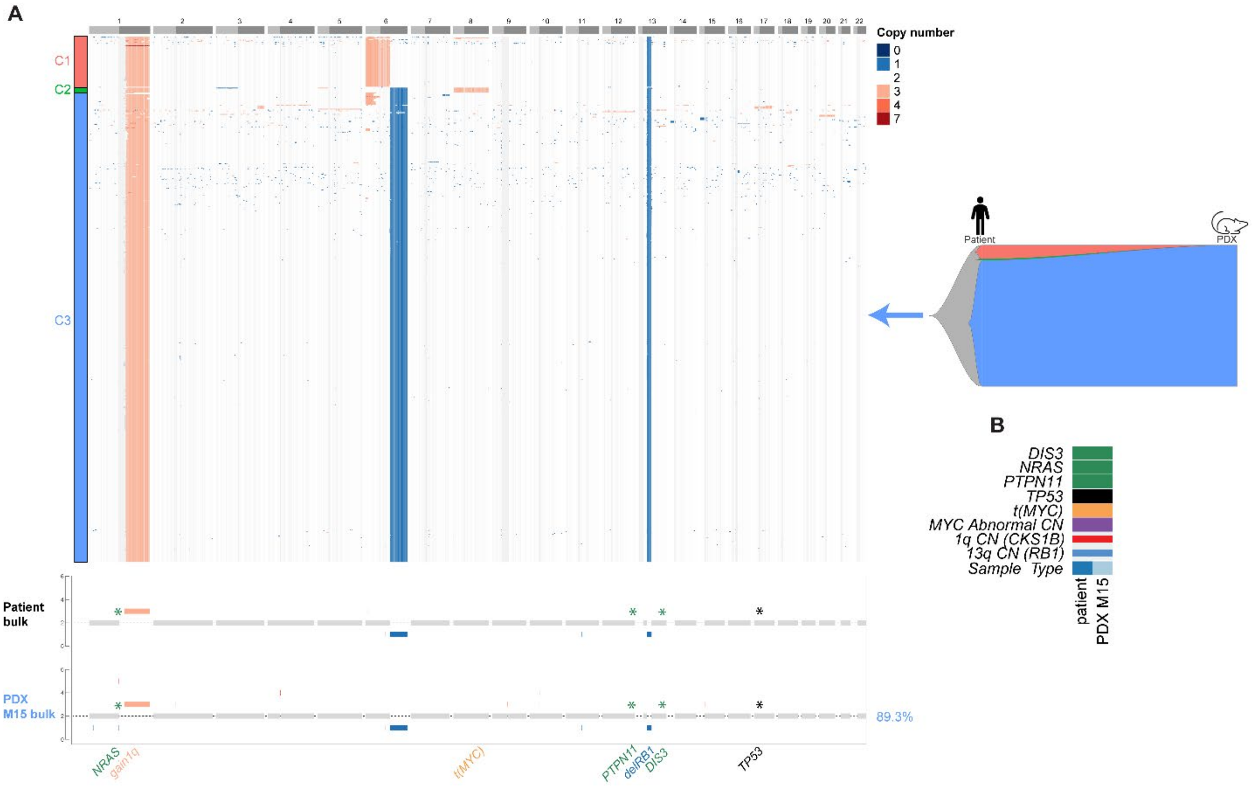

**Extended Data Figure 9. scWGS DLP+ sequencing 0661-900, a hyperdiploid sample.**

**A.** CN defined clones identified by DLP+ scWGS from 566 cells. CN tracks for each sample from bulk WGS are aligned to the scWGS data for comparison. Four main clusters were identified. Fishplots represent the clonal composition of the patient sample at the time of implantation and subsequent selection of clone C2 in the SCID-rab PDX. **B.** MM relevant variants identified by bulk WGS for the patient and corresponding PDXs.

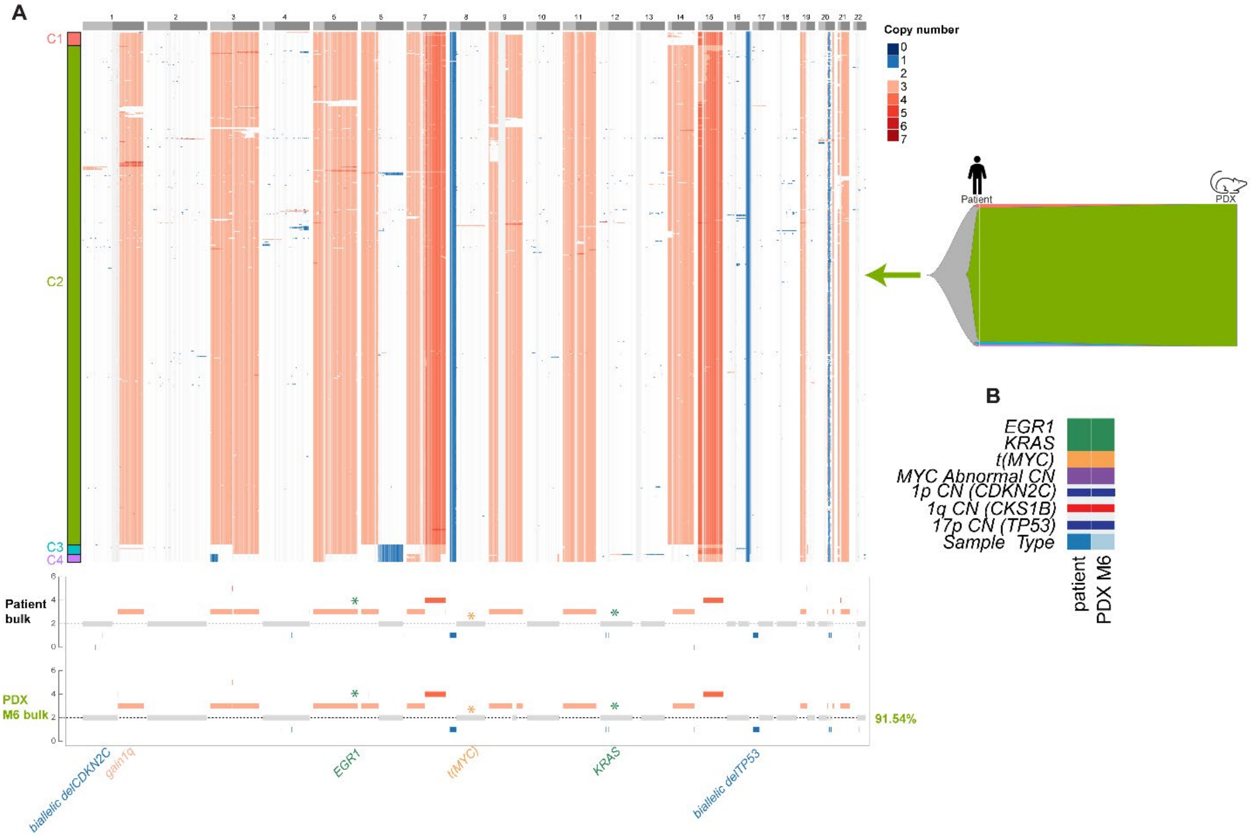
