## Supplementary Data for "Genomic characterization of patient derived xenograft models of multiple myeloma reveals selection of multiple pre-existing high risk clones"

**Running title: PDX models in multiple myeloma**

**Authors**: Nathan J. Becker[1], Enze Liu[1], Riya Sharma[2], Parvathi Sudha[2], Sara Ortega Bertran[1], Oumaima Jaouadi[2], Sreejeta Mondal[1], Nelia Luviano[1], Maurizio Affer[1], Kiran K. Mangalaparthi[3], Akhilesh Pandey[3], Marios Papadimitriou[1], Benjamin T. Diamond[1], David Coffey[1], Abhishek Pandey[1], James Hoffman[1], Gil Hevroni[1], Dickran Kazandjian[1], Kelvin Lee[2], Gareth J Morgan[4], Faith E Davies[4], Attaya Suvannasankha[2], Aneta Mikulasova[5], Ola Landgren[1], Rafat Abonour[1], Brian A. Walker[1]

### Supplementary Data

**Supplementary Figure 1. Determining non-human contamination in PDX cells.**

**A.** To assess host-derived contamination in PDX sequencing libraries, FastQScreen was used to quantify species-specific read alignment. Reads were aligned to the human (hg38), rabbit (Oryctolagus cuniculus, OryCun2.0), and mouse (Mus musculus, GRCm39) reference genomes, with paired patient sequencing libraries included for comparison. Across all P0 PDX samples, the vast majority of reads aligned uniquely to the human reference genome, indicating minimal host-derived contamination. **B.** Where small percentages of reads aligned to a non-human genome they were below the 5% cut-off used in variant callers.

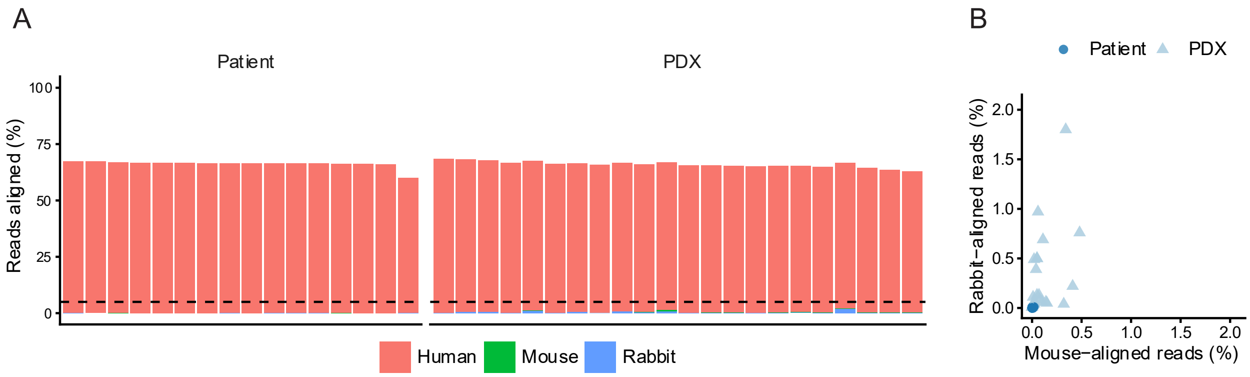

**Supplementary Figure 2. Comparison of variant allele frequencies in the patient and PDX WGS data.** The VAF of coding variants detected in both Patient and PDX paired samples. The VAF of Variants detected in both the patient and PDX were compared using Pearson correlation.

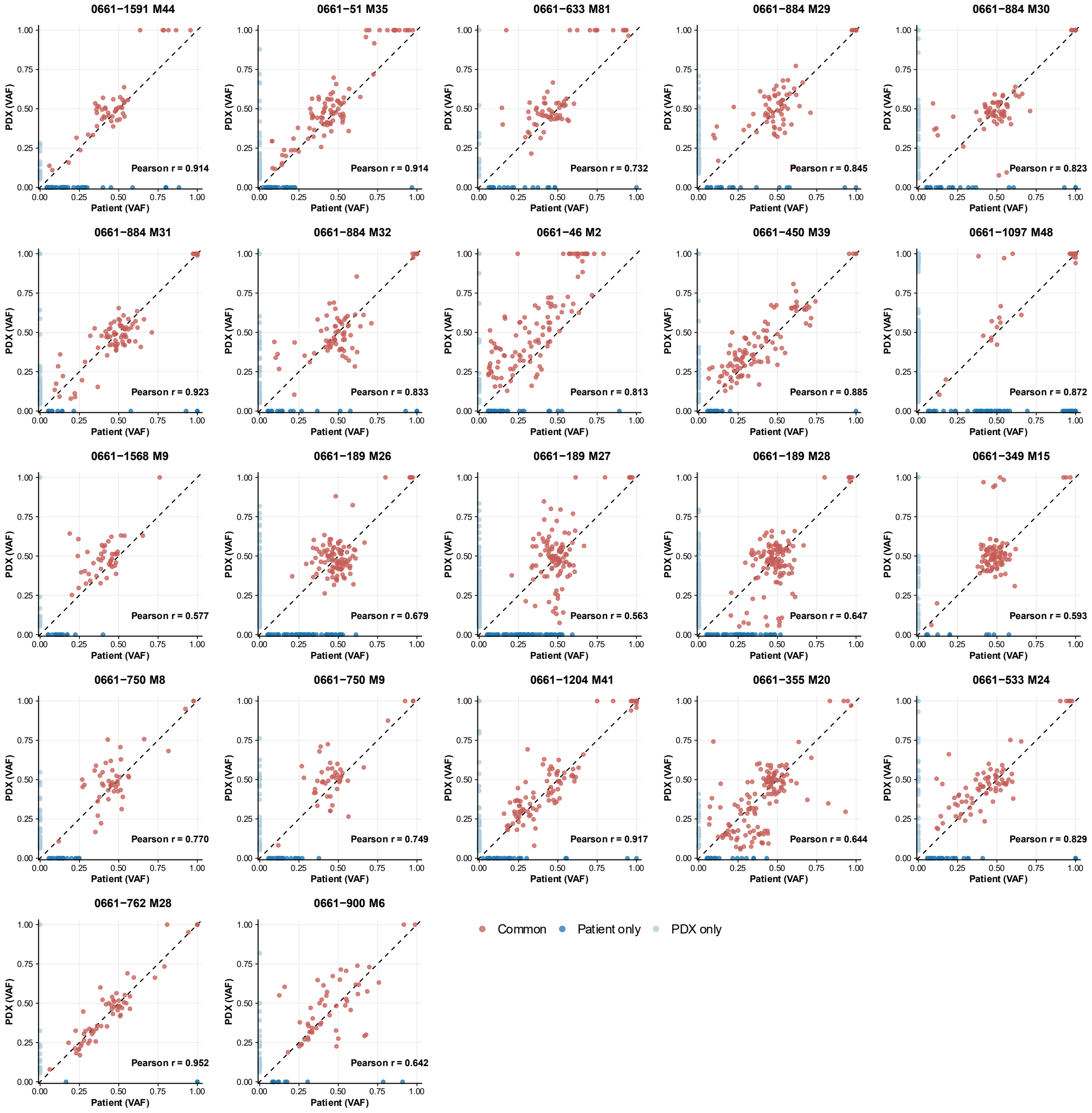

**Supplementary Figure 3.**

ASCAT-NGS profiles for patient and PDX models.

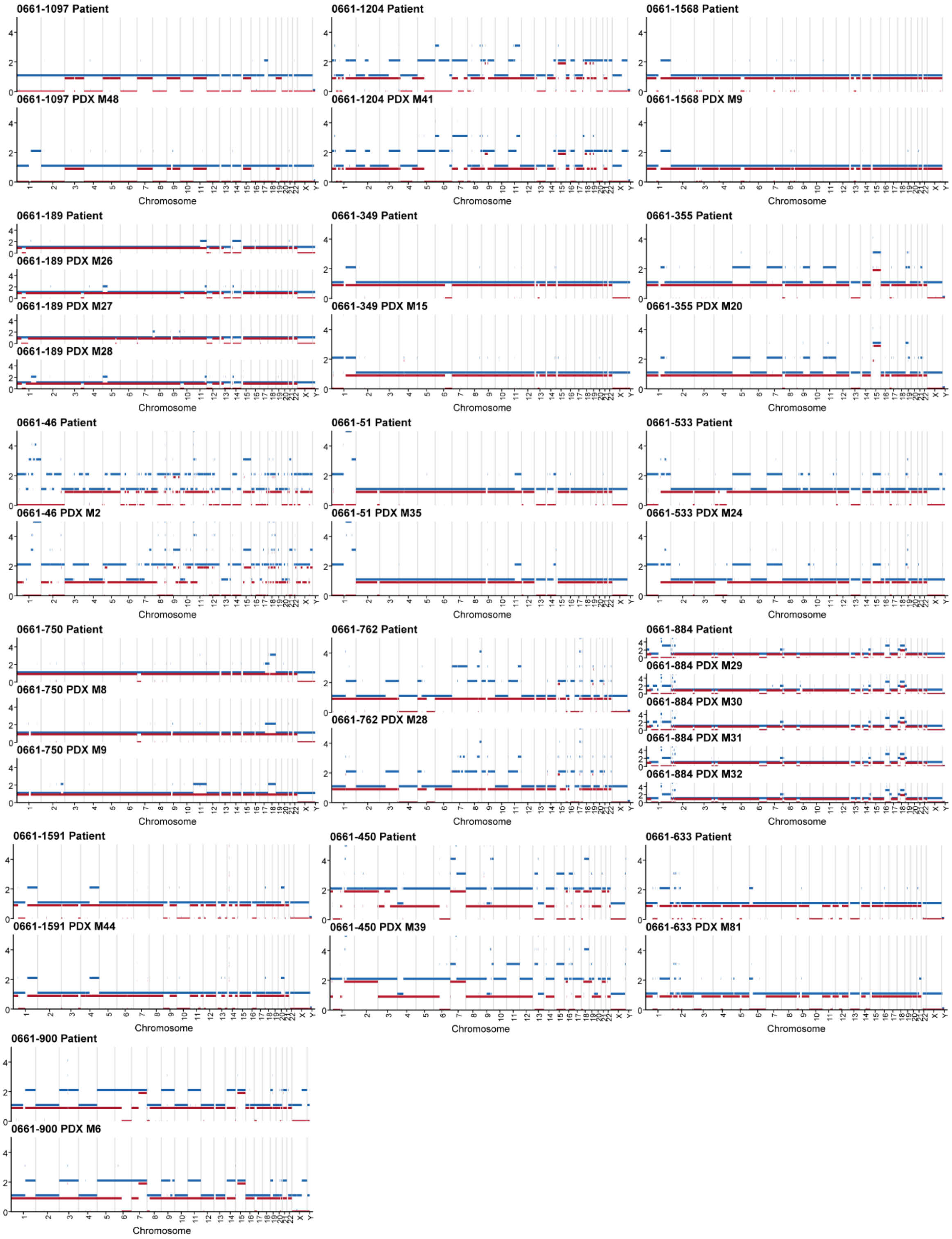

**Supplementary Figure 4. Mutation signature profiles for patient and matched PDXs.** MMSig was used to generate mutation signatures for each patient and PDX sample using all somatic variants from WGS data. The relative contribution of each mutation signature is summarized in **Figure 3**.

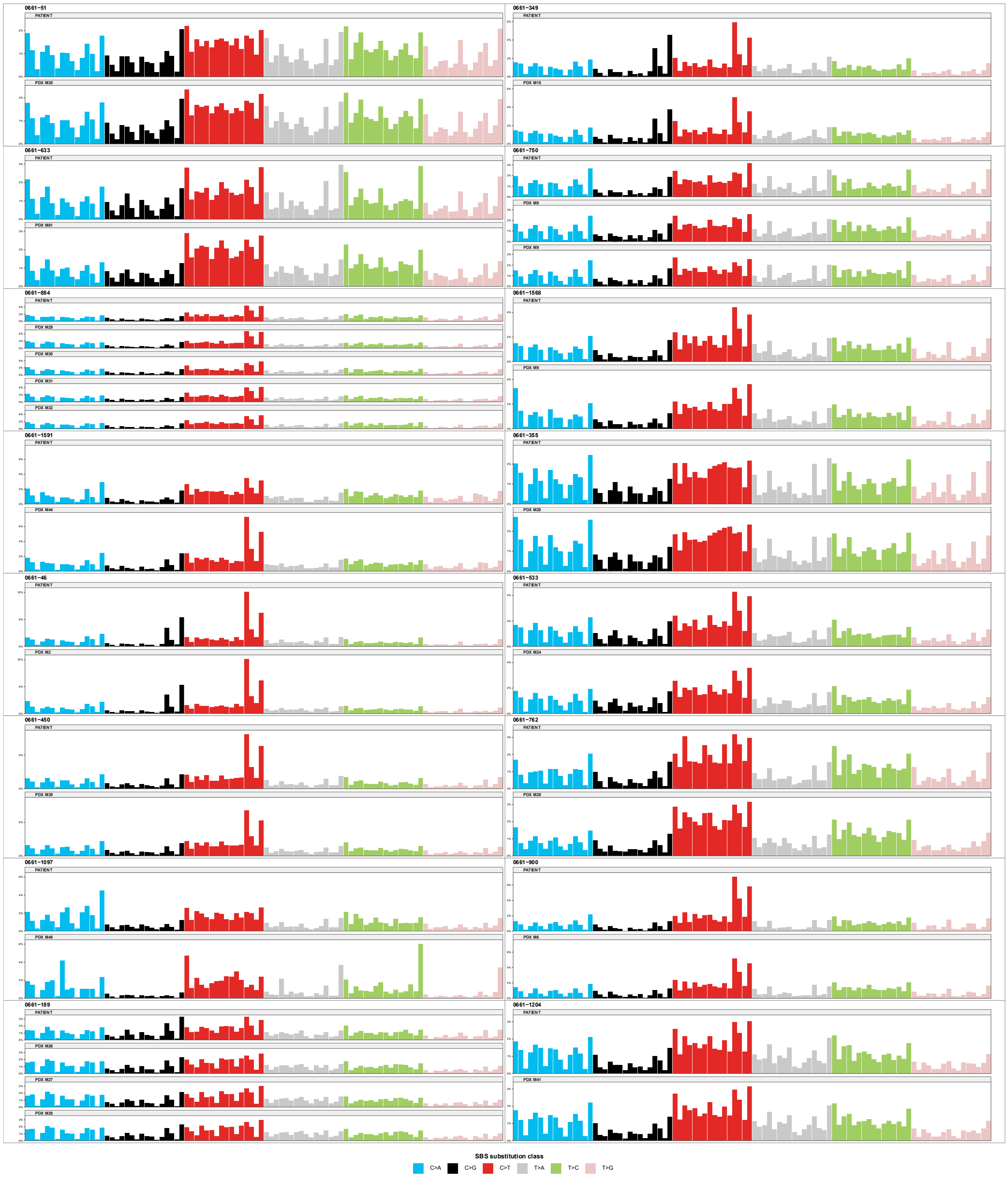

**Supplementary Figure 5. Correlation of RNA expression between patients and PDXs**

RNA expression of patients vs their SCID-rab mice.

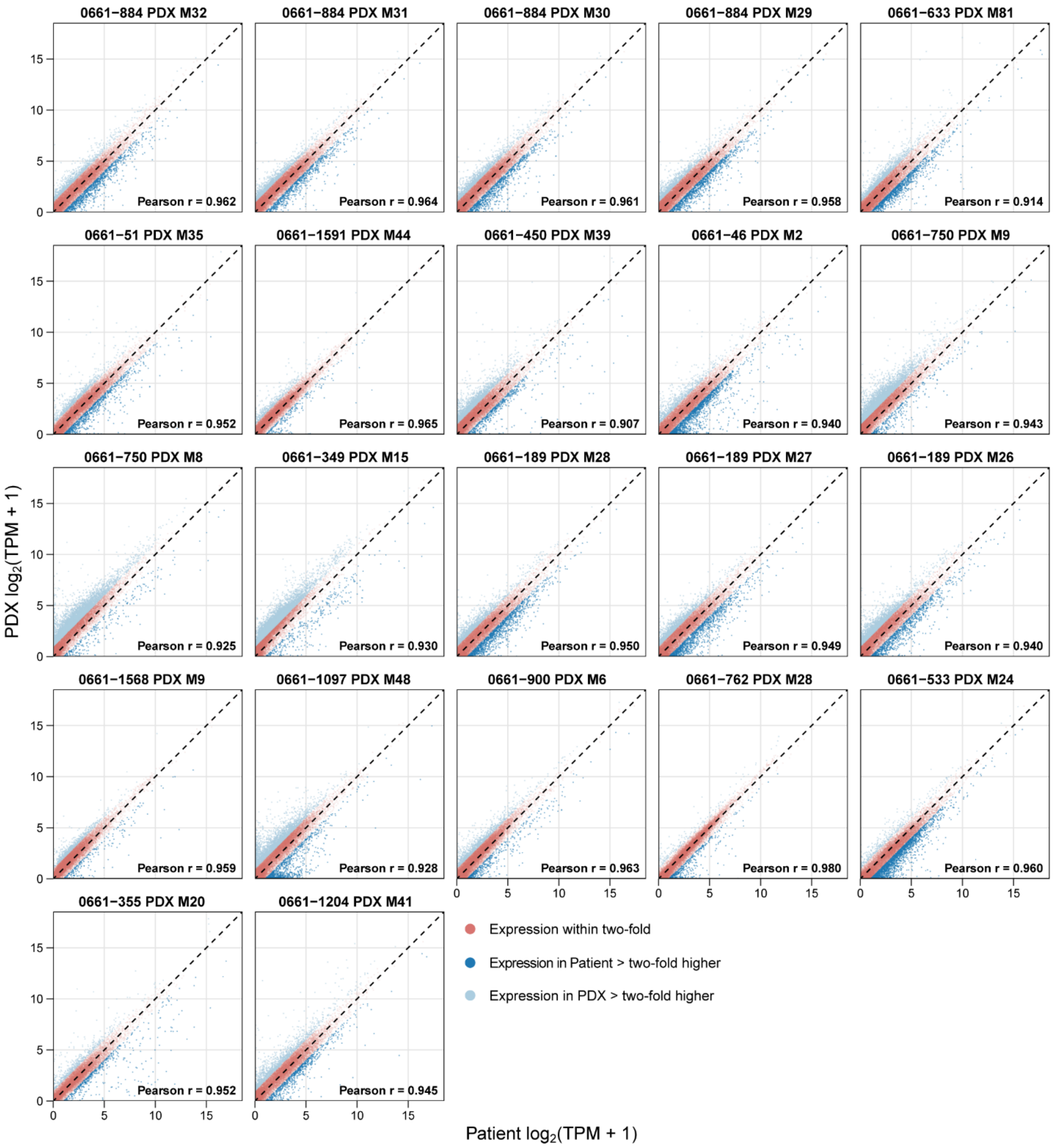

**Supplementary Figure 6. 0661-884 *TP53* focal deletion.**

A 14.4kb focal deletion in *TP53* detected through visualization of WGS BAM files.

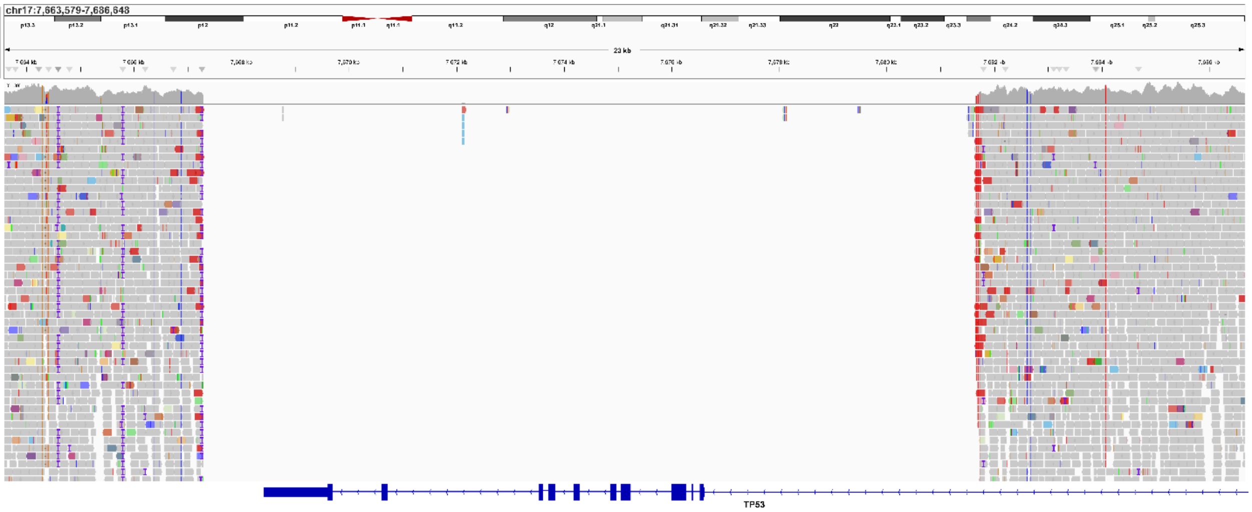

**Supplementary Figure 7. SCID-rab mice maintain a degree of heterogeneity**

VAF clustering of coding mutations by *inferHeterogeneity* was used to estimate heterogeneity SCID-rab PDX mice.

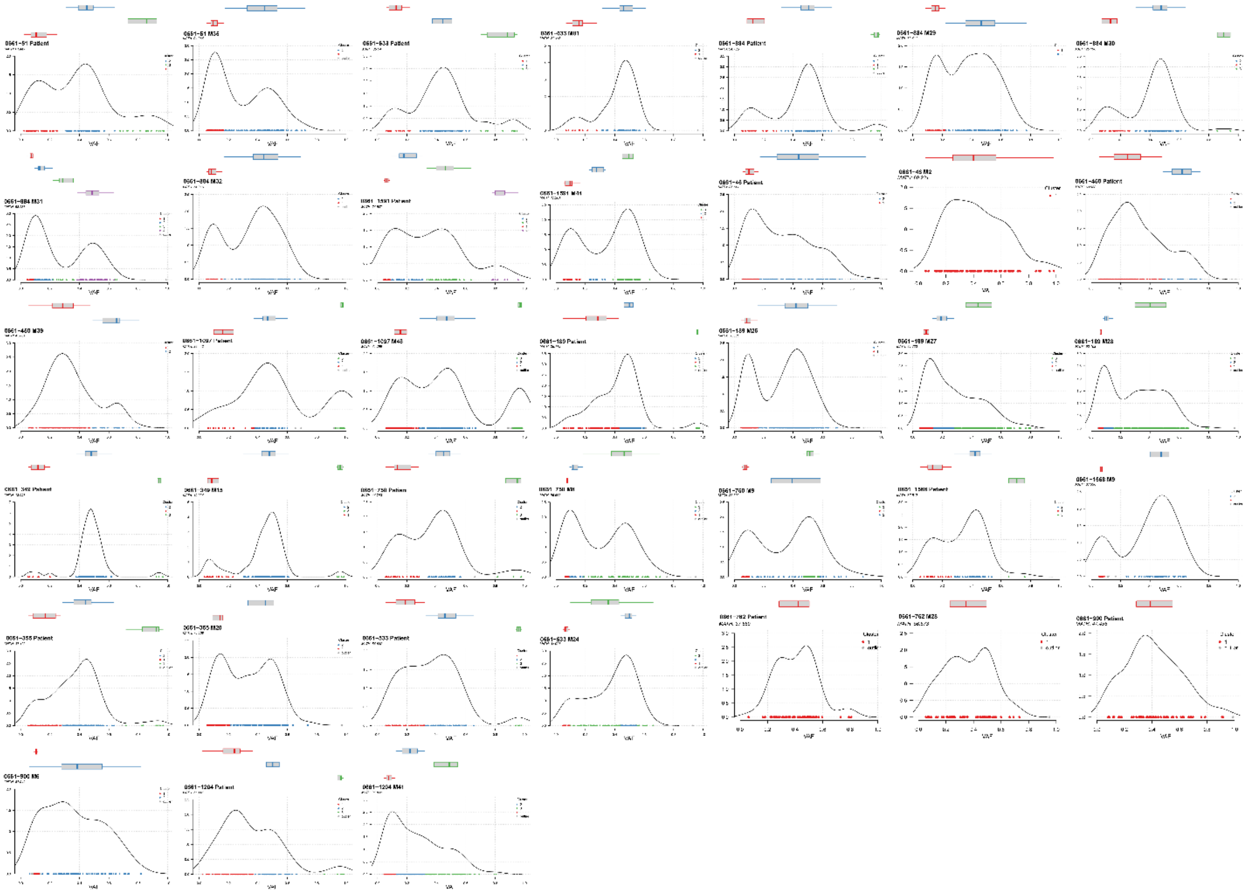

**Supplementary Table 1. Engraftment rates of patient samples.**

| Disease stage | Patients | Patients with recovered cells (%) | Patients with progressive ELISA (%) | Mice | Mice with recovered cells (%) | Mice with progressive ELISA (%) | Mice with both (%) |
|---|---|---|---|---|---|---|---|
| SMM | 7 | 2 (28.6%) | 7 (100.0%) | 9 | 2 (22.2%) | 7 (77.8%) | 2 (22.2%) |
| NDMM | 15 | 4 (26.7%) | 10 (66.7%) | 23 | 5 (21.7%) | 11 (47.8%) | 5 (21.7%) |
| RRMM | 24 | 9 (37.5%) | 19 (79.2%) | 70 | 45 (64.3%) | 54 (77.1%) | 42 (60.0%) |
| sPCL | 8 | 5 (62.5%) | 7 (87.5%) | 38 | 25 (65.8%) | 32 (84.2%) | 24 (63.2%) |
| **Total** | **54** | **20 (37.0%)** | **43 (79.6%)** | **140** | **77 (55.0%)** | **104 (74.3%)** | **73 (52.1%)** |

**Supplementary Table 2. Distribution of skeletal lesions observed in SCID-rab PDX mice**

| Patient ID | Mouse ID | Skeletal regions with clear bone destruction or lesions |
|---|---|---|
| 0661-1097 | 48 | Skull; vertebral column; ribs; pelvis; forelimbs; hindlimbs |
| 0661-1204 | 41 | Forelimbs |
| 0661-1568 | 9 | Skull; vertebral column; pelvis; forelimbs; hindlimbs |
| 0661-1591 | 44 | Skull; vertebral column; pelvis; forelimbs |
| 0661-1591 | 45 | Vertebral column; forelimbs |
| 0661-349 | 47 | Skull; vertebral column; ribs; pelvis; forelimbs; hindlimbs |
| 0661-46 | 2 | Skull; vertebral column; ribs; pelvis; forelimbs; hindlimbs |
| 0661-51 | 49 | Skull |
| 0661-51 | 70 | Hindlimbs |
| 0661-762 | 28 | Vertebral column; forelimbs |
| 0661-884 | 63 | Skull; vertebral column; forelimbs |
| 0661-900 | 56 | Vertebral column; pelvis |

**Supplementary Table 3. scWGS copy number defined subclones**

| Patient ID | Cluster ID | Cells (n) | Cells (%) |
|---|---|---|---|
| 0661-633 | C1 | 10 | 18.87% |
| 0661-633 | C2 | 4 | 7.55% |
| 0661-633 | C3 | 5 | 9.43% |
| 0661-633 | C4 | 1 | 1.89% |
| 0661-633 | C5 | 25 | 47.17% |
| 0661-633 | C6 | 6 | 11.32% |
| 0661-633 | C7 | 2 | 3.77% |
| **0661-633** | **Total** | **53** | **100.00%** |
| 0661-750 | C1 | 3 | 0.31% |
| 0661-750 | C2 | 7 | 0.72% |
| 0661-750 | C3 | 427 | 44.16% |
| 0661-750 | C4 | 3 | 0.31% |
| 0661-750 | C5 | 388 | 40.12% |
| 0661-750 | C6 | 77 | 7.96% |
| 0661-750 | C7 | 5 | 0.52% |
| 0661-750 | C8 | 51 | 5.27% |
| 0661-750 | C9 | 3 | 0.31% |
| 0661-750 | C10 | 3 | 0.31% |
| **0661-750** | **Total** | **967** | **100.00%** |
| 0661-884 | C1 | 6 | 9.09% |
| 0661-884 | C2 | 14 | 21.21% |
| 0661-884 | C3 | 3 | 4.55% |
| 0661-884 | C4 | 5 | 7.58% |
| 0661-884 | C5 | 7 | 10.61% |
| 0661-884 | C6 | 10 | 15.15% |
| 0661-884 | C7 | 2 | 3.03% |
| 0661-884 | C8 | 19 | 28.79% |
| **0661-884** | **Total** | **66** | **100.00%** |
| 0661-900 | C1 | 14 | 2.55% |
| 0661-900 | C2 | 518 | 94.18% |
| 0661-900 | C3 | 10 | 1.82% |
| 0661-900 | C4 | 8 | 1.45% |
| **0661-900** | **Total** | **550** | **100.00%** |
| 0661-349 | C1 | 60 | 9.76% |
| 0661-349 | C2 | 6 | 0.98% |
| 0661-349 | C3 | 549 | 89.27% |
| **0661-349** | **Total** | **615** | **100.00%** |
| 0661-189 | C1 | 7 | 1.08% |
| 0661-189 | C2 | 91 | 13.98% |
| 0661-189 | C3 | 9 | 1.38% |
| 0661-189 | C4 | 3 | 0.46% |
| 0661-189 | C5 | 6 | 0.92% |
| 0661-189 | C6 | 18 | 2.76% |
| 0661-189 | C7 | 517 | 79.42% |
| **0661-189** | **Total** | **651** | **100.00%** |
